# ECT2 peptide sequences outside the YTH domain regulate its m^6^A-RNA binding

**DOI:** 10.1101/2024.08.05.606563

**Authors:** Daphné Seigneurin-Berny, Claire Karczewski, Elise Delaforge, Karen Yaacoub, Celso Gaspar Litholdo, Jean-Jacques Favory, Malene Ringkjøbing Jensen, Cécile Bousquet-Antonelli, André Verdel

**Affiliations:** Université Grenoble Alpes, INSERM U 1209, CNRS UMR 5309, Institut pour l’Avancée des Biosciences, Grenoble, France; Université Grenoble Alpes, CNRS, CEA, Institut de Biologie Structurale, Grenoble, France; CNRS-LGDP-UMR5096, 58 Av. Paul Alduy 66860 Perpignan, France; Université de Perpignan Via Domitia, LGDP-UMR5096, 58 Av. Paul Alduy 66860 Perpignan, FRANCE

**Keywords:** YTH protein, YTH domain, *Arabidopsis thaliana*, m^6^A, RNA binding, IDR, EMSA

## Abstract

The m^6^A epitranscriptomic mark is the most abundant and widespread internal RNA chemical modification, which through the control of RNA acts as an important actor of eukaryote reproduction, growth, morphogenesis and stress response. The main m^6^A readers constitute a super family of proteins with hundreds of members that share a so-called YTH RNA binding domain. The majority of YTH proteins carry no obvious additional domain except for an Intrinsically Disordered Region (IDR). In *Arabidopsis thaliana* IDRs are important for the functional specialization among the different YTH proteins, known as Evolutionarily Conserved *C*-Terminal region, ECT 1 to 12. Here by studying the ECT2 protein and using an *in vitro* biochemical characterization, we show that full length ECT2 and its YTH domain alone have a distinct ability to bind m^6^A, conversely to previously characterized YTH readers. We identify peptide regions outside of ECT2 YTH domain, in the N-terminal IDR, that regulate its binding to m^6^A-methylated RNA. Furthermore, we show that the selectivity of ECT2 binding for m^6^A is enhanced by a high uridine content within its neighboring sequence, where ECT2 N-terminal IDR is believed to contact the target RNA *in vivo*. Finally, we also identify small structural elements, located next to ECT2 YTH domain and conserved in a large set of YTH proteins, that enhance its binding to m^6^A-methylated RNA. We propose from these findings that some of these regulatory regions are not limited to ECT2 or YTH readers of the flowering plants but may be widespread among the eukaryotic YTH readers.

## Introduction

N^6^-methyladenosine (m^6^A) is the most prevalent and evolutionarily conserved of messenger RNA internal chemical modifications in eukaryotes. Present in 1-1.5% of adenosines, this mark contributes to mRNA fate control at all steps of their lives including transcription termination, polyadenylation, splicing, nuclear and cytoplasmic storage, export, stability and translation [1–3]. m^6^A is a mark that is deposited by “writers”, removed by “erasers” and decoded by “readers”. The consensus methylated sequence imposed by the writer methylase is DRACH (D=A, G, U; R=A, G; H=A, C, U). The downstream molecular and cellular effects of m^6^As are conferred by their interaction with reader proteins that are anchored to mRNA by the chemical modification. The YTH-containing protein family constitutes the largest set of m^6^A readers.

The YTH (<u>YT</u>521 <u>H</u>omology) domain is a 100-150 amino acid region, shared by hundreds of eukaryotic proteins, that evolved in two main clades named DF and DC [4–6]. The obtention of 3D structures of YTH domains from yeast [7, 8], human [4, 8, 9], rat [10] and Arabidopsis [11] proteins, further defined the core structure of the YTH domain as a β-barrel formed of three α-helices (α1-3) that stack against six β-strands (β1-6) which form a hydrophobic pocket that tightly and specifically accommodates the m^6^A around three aromatic residues (WW(W/L/Y)), with two tryptophan aromatic rings contacting the methyl group. In several instances, the N- and C-terminal extended YTH domains were used in biophysical studies that revealed the presence of additional upstream and downstream helices, named α0 and α4 [8, 9, 11]. From these studies emerged the general view that YTH domains act as autonomous m^6^A-binding entities, not only necessary but also sufficient to form a functionally competent complex with m^6^A-RNAs. However, the contribution of the α0 and α4 helices, in addition to the core YTH domain, to m^6^A-RNA binding has never been addressed.

Noticeably, the characterized YTH domains of the yeast Mrb1 and Pho92 proteins do not contain these additional α helices, questioning the importance of these two helices acquired during evolution. Furthermore, the early report that the YTH domain of the fly YTHDC1 is unable to form a stable complex alone with m^6^A-RNA *in vitro* [12], suggested that the YTH domain may not necessarily behave as an autonomous m^6^A-binding entity and thus shall require one or more additional peptide sequences, from the same protein or another protein, to stably bind m^6^A-methylated RNA. In addition, in plants, the number of YTH-containing proteins has dramatically expanded, reaching 13 different YTH proteins in the model plant *Arabidopsis thaliana* [5] and up to 39 in the wheat crop *Triticum aestivum* [13]. This large number of YTH proteins capable of binding to the same m^6^A raises the question of possible selectivity in their binding to specific subsets of m^6^A-methylated RNAs, a selectivity that, again, may be brought by one or more additional peptide sequences, from the YTH protein itself or by another protein. In angiosperms (flowering plants), the YTH domains of the DF clade cluster into three subclades tagged as DF-A (comprising Arabidopsis ECT1-4), DF-B (comprising Arabidopsis ECT5, 9-10) and DF-C (comprising Arabidopsis ECT6-8 and 11) [5, 14]. The Arabidopsis ECT2 and 3, which are among the most studied plant YTH proteins, accumulate in dividing cells of organ primordia and exhibit genetic redundancy in the stimulation of stem cell proliferation during organogenesis. Simultaneous loss of ECT2-3, leads to a delay in the emergence of the first true leaves, an aberrant leaf morphology and trichome branching, and a delay in root growth and directionality [5, 15–17]. ECT2 also plays, together with ECT3 and ECT5, a role in the plant’s antiviral defense, and spreading of infection, at least against Alfalfa Mosaic Virus (AMV) [18]. Importantly, both the developmental and antiviral roles of ECT2, directly rely on the ability of its YTH domain to bind m^6^A residues on messenger RNAs (mRNAs) [5, 17, 19]. However, there is no report in the literature that demonstrates the ability of ECT2 YTH domain to be a self-autonomous m^6^A-binding entity.

The regions outside the YTH domain of the DF-type proteins from angiosperms are Intrinsically Disordered Regions (IDRs), and the IDR of Arabidopsis ECT2 protein was recently reported to contribute to its *in vivo* RNA binding activity through iCLIP experiments and RNA co-immunoprecipitation [14, 19]. At the same time, ECT2 IDR has been divided into subdomains that carry distinct *in vivo* molecular functions. Indeed, the N-terminal domain N3 of ECT2 mediates leaf developmental functions by encompassing a conserved Short Linear Motif (SLiM) that mediates a direct physical interaction with a poly(A) Binding Protein [20], while the domain N5 is involved in ECT2 antiviral functions [18]. Nevertheless, a possible relationship between these IDR subdomains and the regulation of ECT2 binding to m^6^A RNA yet remains to be explored.

Here, we uncover that the ECT2 YTH domain alone is not sufficient to bind m^6^A-methylated RNA with a high affinity *in vitro*, compared to ECT2 full-length protein or other YTH domains from Arabidopsis or human readers. We show that the N-terminal IDR of ECT2, as well as structural elements located in the vicinity to the YTH domain (among which the α0 and α4 helices), positively or negatively impact the formation of a stable, high affinity m^6^A-specific complex with RNA. In addition, by studying the importance of the sequence bias of ECT2’s natural m^6^A-methylated RNA targets, we demonstrate that the uridine content, surrounding the m^6^A DRACH motif, further inclines ECT2’s binding towards the m^6^A-methylated form of its target RNAs. This work, combined with the recently published *in vivo* studies on ECT2, sheds light on how regulation and selectivity of ECT2 binding to m^6^A RNA might be achieved. Furthermore, based on these findings and Alphafold structural modelling of other YTH proteins, we propose that this m^6^A-binding regulation and selectivity might apply in a similar way to other YTH proteins from plants and other eukaryotes.

## Material & methods

### Construction of ECT2 truncated and deletion mutants, YTH domains of CPSF30L and hYTHDC1

To construct all the ECT2 truncated mutants described in this study, we used the recombinant plasmids p817 (containing ECT2 coding DNA sequence (CDS)) and p883 (a mutated version of ECT2 CDS in which the three tryptophans at positions 464, 521 and 526 in the YTH domain were substituted to alanines) as previously described [5]. The constructs ECT2-108/667, 204/667, 303/667, 376/667, 424/610 and 444/580 (respectively named ECT2-108, -204, -303, -376, -424 and -444) were obtained by PCR amplification using primers listed in Suppl. Table S1. Primers were designed to introduce a *Bam*HI restriction site in 5’ and a *Xho*I restriction site in 3’ of the DNA fragment. The amplified fragments were digested with *Bam*HI and *Xho*I, and inserted into the expression vector pETM11-SUMO3-eGFP (from EMBL) previously digested with *BamH*I and *Xho*I (this digestion removes eGFP CDS). The resulting expression plasmids allow the expression of N-terminal His-SUMO tagged versions of the ECT2 protein. For the production of His-tagged proteins, the truncated forms ECT2-303/667 and ECT2-424/610 were obtained after PCR amplification with primers containing *Nde*I restriction site in 5’ and *BamH*I restriction site in 3’ of the DNA fragment (Suppl. Table S1). After digestion with *Nde*I and *BamH*I, the inserts were ligated into the expression vector pNEA-3H previously digested with *Nde*I and *BamH*I. For the production of proteins with a cleavable His-SUMO tag, the pETM11SS vector (from Ramesh Pillai lab, Geneva University) was used and the cDNA inserted between *BamH*I and *Xho*I restriction sites. This vector allows the production of recombinant proteins with a His-Strep-SUMO tag in N-terminal that it is cleavable with a TEV protease.

The deletion mutants Δ376-386, Δ419-427 and Δ591-627 were obtained from the pETM11-SUMO3_ECT2 recombinant plasmid by site-directed mutagenesis (Q5 Site-Directed Mutagenesis Kit, BioLabs) using the primers listed in Suppl. Table S1.

The construct CPSF30L 220-400 and YTHDC1 345-507 were obtained by PCR amplification from respectively pBSK plasmid containing the Arabidopsis CDS of CPSF30L (ordered from Biomatik) and a pCI-neo-hYTHDC1 plasmid already produced [21] using primers listed in Suppl. Table S1. Primers were designed to introduce a *BamH*I restriction site in 5’ and a *Xho*I restriction site in 3’ of the DNA fragment. The amplified fragments were digested with *BamH*I and *Xho*I and inserted into the expression vector pETM11-SUMO3-eGFP previously digested with *BamH*I and *Xho*I. All the constructs obtained were validated by sequencing.

### Production in E. coli and purification of the recombinant proteins

Recombinant plasmids were transformed into *Escherichia coli* strain BL-21(DE3) competent cells. The *E. coli* cells were grown at 37°C to an OD600 of 0.6, and recombinant protein expression was then induced with 1 mM IPTG overnight (16h) at 20°C. The pellet from 500 mL culture was collected by centrifugation 15 min at 2220 g at 4°C and resuspended in 20 mL of Lysis buffer (20 mM Tris-HCl, pH 7.5, 150 mM NaCl, 10 mM imidazole, 1 mM PMSF) containing 1% Triton X-100, and incubate 10 min on ice. The suspension was sonicated for 2 min (30” on, 30” off) using a probe sonicator (Active motif), at 85% amplitude. The sample was centrifuged at 20800 g for 20 min at 4°C, and the supernatant collected. The supernatant was then incubated with Ni-NTA agarose (Qiagen, 500 µL of agarose for 500 mL of bacteria culture) that had been preequilibrated in the lysis buffer (without Triton). After 1h of incubation at 4°C on a rotating shaker, the agarose/sample mixture was loaded onto a column. The resin was then washed with 4 volumes of Lysis buffer, 6 volumes of Wash buffer 1 (20 mM Tris-HCl, pH 7.5, 150 mM NaCl, 40 mM imidazole), 1 volume of Wash buffer 2 (20 mM Tris-HCl, pH 7.5, 500 mM NaCl, 40 mM imidazole) and 1 more volume of Wash buffer 1. The bound proteins were eluted with 4 volumes of Elution buffer (20 mM Tris-HCl, pH 7.5, 300 mM NaCl, 200 mM imidazole) and collected by 500 µL fractions. The purified proteins were desalted against 20 mM Tris-HCl, pH 7.5, 150 mM NaCl (PD-10 column, Merck), concentrated using centrifugal filter unit, and stored at -80°C after addition of 5% glycerol. Purified recombinant proteins were quantified using the BIORAD protein assay reagent [22] before SDS-PAGE analyses and EMSA assays.

For cleavage reactions, 600 μg of fusion protein were used. Reactions were conducted in 300 µL with the following reagents added to the final concentrations of 1 mM DTT, 0.5 mM EDTA in 20 mM Tris-HCl, pH 7.5, 150 mM NaCl, and 10 µL of commercial TEV (Biolabs). After incubation overnight at 4°C, the suspension was diluted 4 times and incubated for 1 hour at 4°C with 200 µL of Ni-NTA agarose (Qiagen) preequilibrated in the lysis buffer. The flow through was recovered, concentrated using centrifugal filter unit, and stored at -80°C after addition of 5% glycerol. For NMR analyses, cleavage was done from 10 mg of tagged proteins, the cleaved proteins were desalted against NaCl 50 mM, sodium phosphate 50 mM pH7, concentrated and flash-frozen without glycerol before storage at -80°C.

### NMR measurements

The YTH-ECT2 buffer contained 50 mM sodium phosphate pH 7, 50 mM NaCl and 10% (v/v) D2O. A one-dimensional ^1^H NMR spectrum of YTH-ECT2 at 270 µM was recorded using a Bruker spectrometer operating at a ^1^H frequency of 600 MHz, equipped with a cryoprobe. The spectrum was processed using Topspin 4.1.4.

### Analysis of in vitro RNA binding by electrophoretic mobility shift assay (EMSA)

RNA probes were labelled with the 6-FAM fluorophore and synthesized by IBA Lifesciences (for probes I), and LGC Biosearch technologies for the other ones. The probe sequences are listed in Suppl. Table S1. The final RNA probe concentration was 10 nM, and the concentration of the purified recombinant proteins ranged from 0 to 500 nM in most assays (in some case up to 1000 nM). The probe was heated to 65°C for 10 min, and then slowly cooled down to room temperature. The purified recombinant proteins were diluted to concentration series (10, 5, 2.5, 2, 1, 0.5, and 0.25 µM in most case) in binding buffer (10 mM HEPES, pH 8.0, 50 mM KCl, 1mM EDTA, 0.05% Triton-X-100, 5% glycerol, 10 µg/mL Salmon DNA, 1 mM DTT and 40 U/mL RNasin). For the reaction, 4 µL of RNA probe (at 25 nM) and 2 µl of protein dilution were added in a final reaction volume of 10 µL. After 20 min incubation on ice, 9 µL of the mixture was loaded onto a native 8% acrylamide TBE gel, and run at 4°C for 5 min at 100V and then 90 min at 80V. Fluorescent signals were detected using a ChemiDocMP system (BioRad) and quantification done with the ImageLab software (Biorad). The apparent Kds (dissociation constant) were calculated with nonlinear curve fitting using the PRISM software (nonlinear regression method).

### 3D structure prediction

Alphafold was used to obtain the predicted 3D structure of Arabidopsis ECT2, ECT5, ECT8, and CPSF30L proteins, human YTH proteins (YTHDF1-3, DC1-2), Drosophila and rat YTHDC1 proteins and yeast Mrb1, Pho92 YTH proteins (https://alphafold.ebi.ac.uk/; [23]). The predicted structures were then analyzed using the Chimera software.

## Results

### ECT2 YTH domain binding to m^6^A-methylated RNA does not form a stable complex in vitro

To characterize ECT2 binding to m^6^A-methylated RNA, we performed Electrophoretic Mobility Shift Assays (EMSAs) using the full-length protein or an extended ECT2 YTH domain (ECT2-424) encompassing the YTH domain (as defined in Uniprot, Suppl. Figure 1A, Suppl. Table S2) plus 20 and 30 amino acid extensions, respectively, at its N- and C*-*termini (Figure 1A). As shown previously [17], the full-length ECT2 (Fl) forms a slow migrating complex with m^6^A-methylated RNA probe while no complex formation is observed with an identical non-methylated probe (Figure 1B). Noticeably, the RNA-ECT2 Fl complex migrates as a doublet suggesting the possible formation of a dimeric complex. Both the binding of ECT2 to m^6^A-RNA and the formation of a higher order complex require an intact aromatic pocket within the YTH domain, as substitution of the three tryptophans with alanines blocks the formation of any RNA-protein complex (Suppl. Figure 1B, [17]). In contrast, the extended YTH domain ECT2-424 is unable to form a slow-migrating RNA-protein complex despite a clear reduction in the amount of the free m^6^A probe, as shown by signal intensity decrease (Figure 1C, left panel; Suppl. Figure 1C). Such decrease in the free probe levels is not observed when using the non-methylated version of the RNA probe (Figure 1C, right panel). This suggests that m^6^A-dependent binding of ECT2-424 to RNA forms an unstable complex. The one-dimension ^1^H NMR spectrum of the ECT2-424 construct shows that this is not because of folding issues of the recombinant protein fragment (Figure 1D). Importantly, the extended YTH domains of human YTHDC1 and Arabidopsis CPSF30-L do form a complex with the m^6^A-RNA probe that resists gel electrophoresis migration, as published [9, 11] (Figure 1E, Suppl. Figure 1A, 1D). To eliminate the possibility of a negative impact of amino acids surrounding the YTH domain in the case ECT2-424 construct, we also performed EMSA with the core ECT2 YTH domain (ECT2-444, aa 444-580, without the predicted α0 and α4 helices). No binding was observed with this YTH construct (Suppl. Figures 1E, 1F). Taken together, these results show that, unlike most of the characterized YTH domains, the YTH domain of ECT2 alone or together with the surrounding α0 and α4 helices does not form a complex with m^6^A-methylated RNA that resists gel migration, and that one or more domains of ECT2 located further outside of the YTH are required for such stabilization.

**Figure 1.**
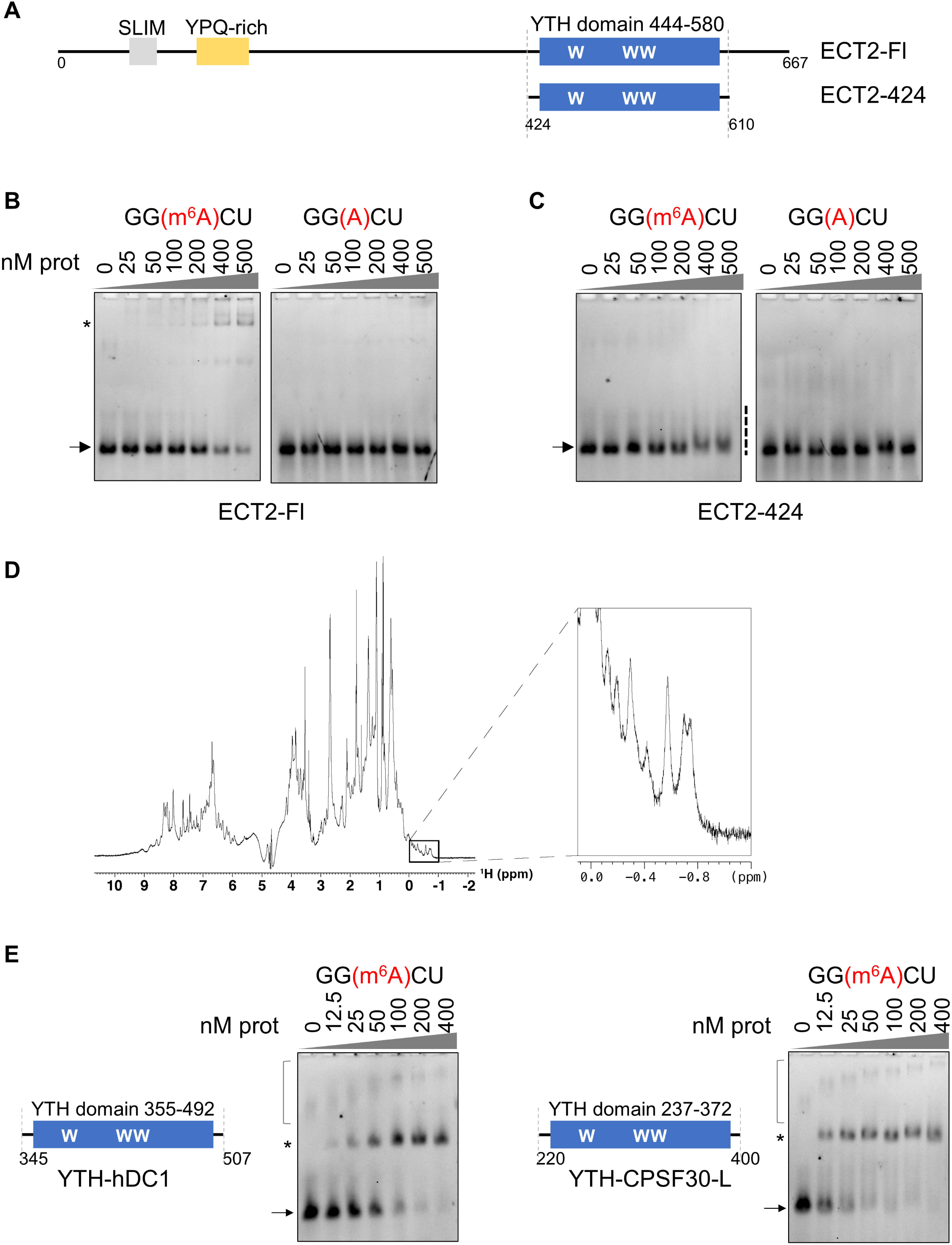
ECT2 YTH domain binding to m^6^A-methylated RNA does not form a stable complex *in vitro*. **A.** Graphical representation of the full-length ECT2 protein (ECT2-Fl) and its extended YTH domain “ECT2-424”, aa 424-610, with the core YTH spanning aa 444 to 580 (in blue) as described in Uniprot. The N-terminal part of ECT2 contains a Short Linear Motif (SLiM enriched in tyrosines, in grey) [20] and a YPQ-rich motif (in yellow) [5]. **B & C.** EMSA gels representing the binding capacity of ECT2-Fl (B) and ECT2-424 (C) to methylated and non-methylated RNA probes. Purified non-tagged proteins (from 0 to 500 nM) were incubated with 10 nM of Fam-labelled RNA probes and loaded onto a native acrylamide gel. Detection of the free and bound probes was done using ChemiDocMP System (BioRad). RNA Probe I: 5’-AUGGGCCGUUCAUCUGCUAAAA(GGXCU)GCUUUUGGGGCUUGU-3’, X = m^6^A/A. Only 5 residues are shown on the figure. The arrows indicate the free probe, the stars the probe/protein complexes, and the dashed line, the smear in the case of the ECT2-424 construct. **D**. ^1^H NMR spectrum of ECT2-424 showing that the recombinant protein is folded in solution. Amide resonances (7-9.5 ppm) are widely dispersed and several side-chain methyl resonances are found below 0 ppm, both characteristic of folded proteins [27]. **E**. EMSA gels performed with extended YTH domains of proteins from the DC family. The purified His-SUMO tagged domains of the human YTHDC1 (hDC1) protein (aa 345 to 507) and the Arabidopsis CPSF30-L protein (aa 220 to 400) were incubated with 10 nM of methylated RNA probe I. The protein concentration range was from 0 to 400 nM. The arrows indicate the free probe, the stars the probe/protein complexes. Non-specific signal is shown by the square bracket.

### The N-terminal IDR of ECT2 modulates the binding of its YTH to m^6^A

The N-terminal part of ECT2 before its YTH domain forms a long Intrinsically Disordered Region (IDR, Figure 2A), which is also the case for most of the readers from the YTHDF family of flowering plants [14]. IDRs have the potential to govern the specificity and affinity of structured RNA binding modules [24]. In addition, i*n vivo* data from iCLIP experiments have indicated that, in addition to its YTH domain, the IDR of ECT2 also contacts ECT2 target mRNAs, suggesting that the IDR-dependent binding may influence ECT2 YTH domain binding to m^6^A [19]. Based on the sequence comparison of YTHDFA proteins from angiosperms (Suppl. Figure 2A, Suppl. Table S3) combined with the prediction of ordered and disordered regions (Figure 2A), we expressed and purified four truncated versions of ECT2, cutting from its N-terminus towards the YTH domain, approximately every 100 amino acids and outside potentially small ordered peptides, successively eliminating the different unstructured regions of the IDR (Figure 2B, Suppl. Figures 2B, 2C). We also prepared for each of these four truncated versions (namely, ECT2-108, ECT2-204, ECT2-303 and ECT2-376), a mutant version with the triple Trp to Ala substitution that inactivates the binding of the YTH pocket to m^6^A.

**Figure 2.**
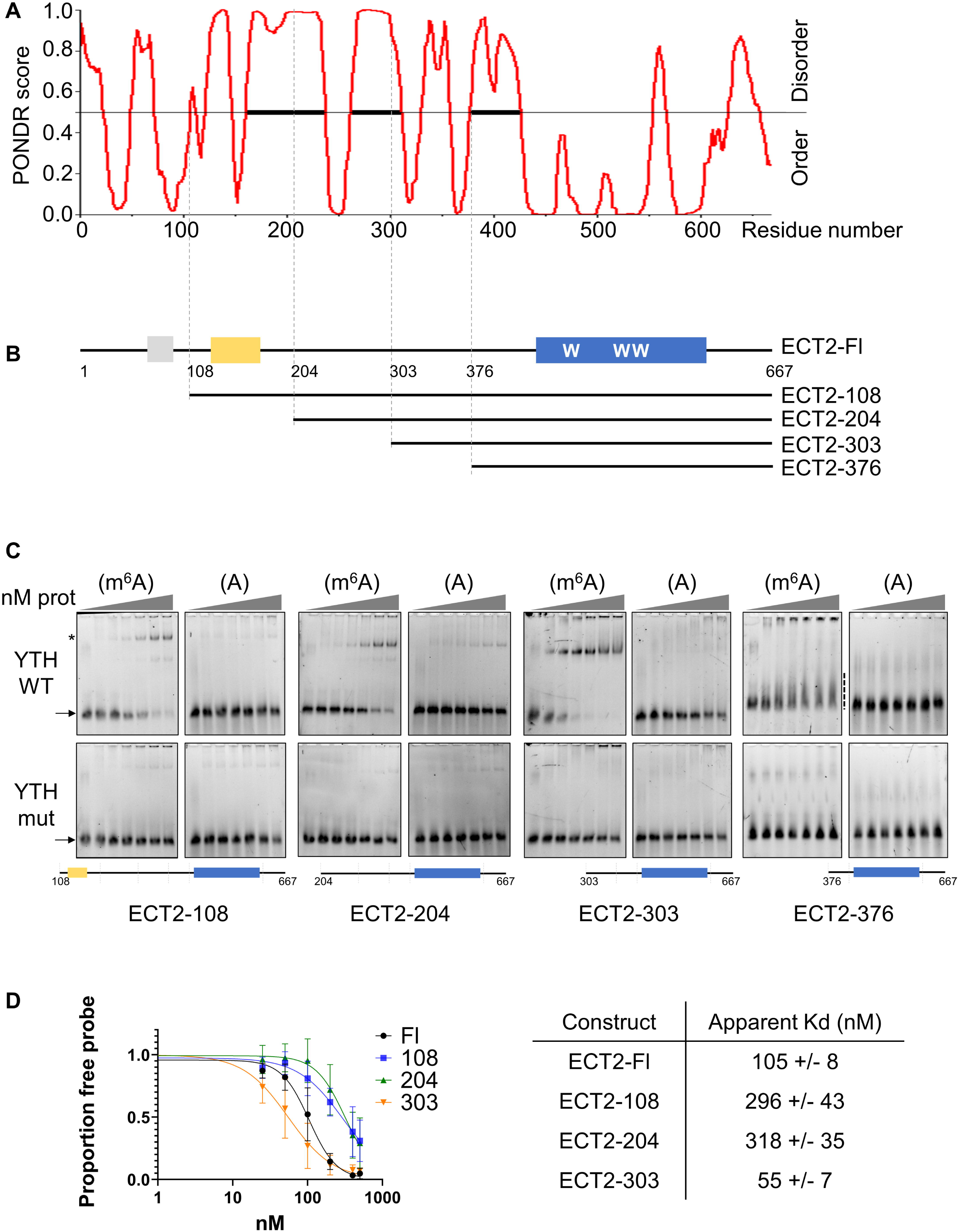
ECT2 N-terminal IDR modulates YTH binding to m^6^A. **A.** Diagram showing the distribution of disordered and ordered regions along the full-length ECT2 protein using the PONDR® VL-XT software [28]. **B.** Graphical representation of ECT2. The SLiM and YPQ-rich regions are shown respectively in grey and yellow, the YTH domain in blue (with the tryptophans required for m^6^A recognition). The truncated forms of ECT2 used for EMSA assays are presented below the full-length sequence of ECT2. The drawing of ECT2 in B can be directly compared to the diagram in A, as identical scales used were. **C.** EMSA gels performed with various concentrations of the purified His-SUMO tagged truncated forms of ECT2 (from 0 to 500 nM; 0-25-50-100-200-400-500 nM) in the presence of 10 nM of methylated (m^6^A) or non-methylated (A) 42 nt RNA probe I. For each construct, a mutated version of the YTH domain (the three tryptophans required for m^6^A binding mutated into alanines, “YTH mut”) was analyzed. Drawings of each construct are shown under the EMSA gels. The arrows indicate the free probe, the star the probe/protein complexes, and the dashed line, the smear in the case of non-stable complex (for the ECT2-376). **D.** Signals of the free probe were quantified with the ImageLab software (Biorad). Graphs show the mean and standard deviation of at least three independent experiments. The curve fittings and the apparent Kds were obtained using the PRISM software.

Interestingly, EMSAs show that ECT2-108, ECT2-204 and ECT2-303 recombinant proteins all bind the RNA methylated probe, while ECT2-376 does not (Figure 2C). As seen for ECT2-424, EMSAs of ECT2-376 show a decrease in the free probe signal and the appearance of a smear above, supporting again that the RNA-protein complex may form but that it is not stable enough to resist gel electrophoresis. This indicates that the ECT2 region encompassing amino acids 303 to 375 is necessary to stabilize the complex formed by ECT2 YTH domain binding to the m^6^A-methylated RNA. As expected, EMSAs conducted with the non-methylated version of the RNA probe and/or the tryptophan point mutant versions of the YTH pocket, do not show a slow migrating complex nor reduction in the levels of free probes for the ECT2-108 and ECT2-303 constructs (Figure 2C, lower panels).

We then quantified the RNA binding affinities of full-length ECT2 and its truncated forms (Figure 2D). Our results show that the ECT2-303 construct possesses the highest apparent affinity and that domains located at its N-terminal side influence either negatively or positively ECT2 YTH domain affinity for m^6^A-methylated RNA. Indeed, addition of the 204-302 region to ECT2-303 construct leads to a 5-fold decrease in the complex dissociation constant, Kd. Further addition of the 108 to 203 region did not affect the affinity compared to that of ECT2-204 construct, whereas the final addition of the domain 1-107 improved the affinity by approximately 3-fold, as compared to ECT2-108 or ECT2-204 constructs. Noticeably, similar differences in binding properties were observed independently of the presence of the tag used to purify the recombinant proteins (Suppl. Figures 2D-G). We also observed that although ECT2-108, ECT2-204 and ECT2-303 constructs stably bind methylated RNA, the RNA-protein complex doublet observed with the full-length ECT2 protein is no longer formed (Figure 2C, upper panels). This suggests that the region encompassing the first 108 amino-acids may also contain an ECT2 dimerization domain. Altogether, these results show that the domain “303-375” of ECT2, which contains several conserved amino acids shared among YTHDFA proteins in angiosperm, is required to stabilize the complex formed by ECT2 YTH domain binding to m^6^A-methylated RNA and that the domains “1-108” and “204-302” may play a regulatory role by controlling the binding affinity.

### Binding of ECT2 to the m^6^A-methylated DRACH motif is enhanced by the surrounding RNA sequence

Biochemical characterizations of YTH domains binding to RNA were conducted with short RNA probes, from 5 to 11 nucleotides (*i.e*. 5, 9 or 11 nt long probes for human YTHDC1, [8, 25]; 10 nt for Arabidopsis CPSF30-L [11]) supporting the fact that the DRACH motif can be sufficient for the binding of YTH domains to m^6^A-methylated RNA. Furthermore, the binding of YTH domains to m^6^A-methylated DRACH motif has been reported to implicate at most ribonucleotides spanning positions +2 to -2 [8] relative to the methylated adenosine, thus within the DRACH motif itself. However, this only relies on assays conducted with self-sufficient YTH domains, that form *per se* a stable RNA-protein complex *in vitro*. Our EMSAs from Figures 1 and 2 show that the YTH domain from ECT2 is necessary but not sufficient to form a complex with m^6^A-methylated RNA that resists to gel migration, and that peptide sequences located N-terminally to the YTH domain are required for stabilization of the complex. Importantly, *in vivo,* the IDR of ECT2 is directly contacting RNA sequences located near to the DRACH motif [19, 20]. We therefore assessed *in vitro* whether the length of the RNA plays a role in the formation or stability of a m^6^A-dependent RNA-protein complex. Figures 1 and 2 show EMSAs conducted with a 42 nt-long RNA probe (probe I) that was reported to form a high-affinity complex with the full-length ECT2 protein [17]. Here, we designed a shorter 11 nt-long RNA probe (probe II) containing the DRACH (GGACU) motif either methylated or not (Figure 3A). We then performed the EMSAs with the probe II and the full-length and truncated forms of ECT2 (Figure 3B). Strikingly, all protein constructs lost their ability to form a stable complex with m^6^A-RNA when incubated with probe II. Nevertheless, for each construct, the signal of the free methylated short probe decreased (probe II m^6^A, Figure 3B, Suppl. Figure 3). These findings show that ECT2 YTH domain is self-sufficient for the recognition of m^6^A methylated DRACH while the formation of RNA-protein complexes that resist to the gel migration also requires extra-RNA sequences near the DRACH motif.

**Figure 3.**
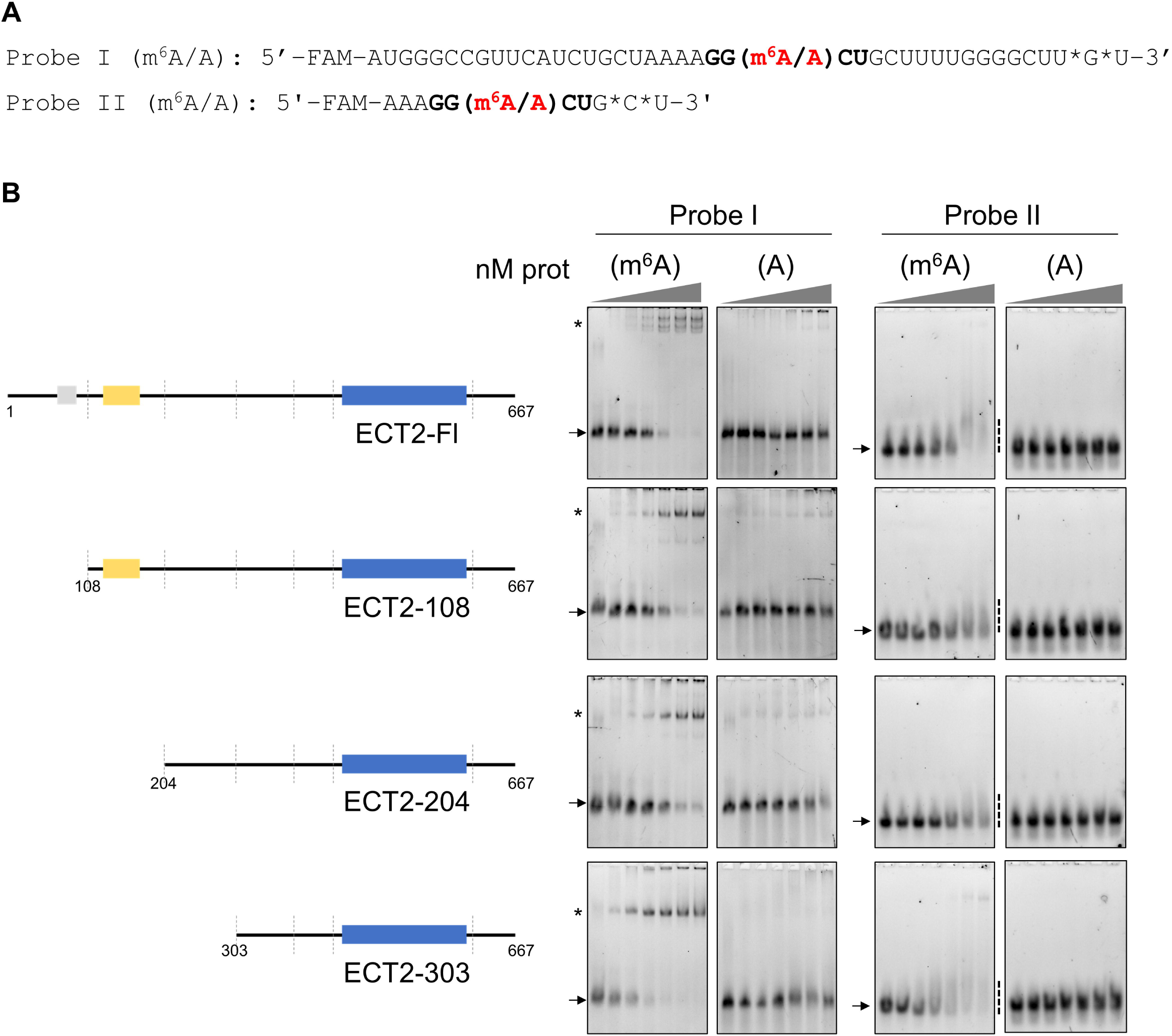
Binding of ECT2 to the m^6^A-methylated DRACH motif is enhanced by the surrounding RNA. **A.** Sequence of the RNA probes I and II used for the EMSA assays. The asterisks indicate thiol-protected bases to avoid 3’-degradation of the RNA. The DRACH motif is shown in bold with the methylated or non-methylated adenosine. Probe I (m^6^A/A) was the one used in the study by Wei and collaborators [17] and in assays of Figures 1 and 2. **B.** EMSA gels obtained with His-SUMO tagged full-length and truncated ECT2 proteins. In each assay, 10 nM of methylated (m^6^A) or non-methylated (A) probes were incubated with increasing concentrations of purified proteins (from 0 to 500 nM). A schematic representation of the various constructs is shown on the left hand-side. The arrows indicate the free probe, the stars the probe/protein complexes, and the dashed line, the smear in the case of unstable complex with the probe II.

### A U-rich content in the RNA sequence around the DRACH motif reduces ECT2 m^6^A-independent binding to RNA

To further characterize the importance of the RNA sequence around the DRACH motif, we performed EMSAs with long RNA probes of equal lengths but of distinct sequence contents. We based their design on results from *in vivo* iCLIP experiments indicating that various U-rich motifs contacted by ECT2’s N-terminal IDR may promote ECT2 binding to m^6^A-methylated mRNAs [19]. We designed two new 42 nt-long probes in which either every uridine was replaced by adenosine, or the U-rich motifs identified by Arribas-Hernandez and colleagues were added (respectively probe III and IV, Figure 4A). For each probe, methylated and non-methylated versions were also tested (Figure 4A). All of the ECT2 constructs containing the first 302 amino-acids of ECT2 form RNA-protein complexes with every methylated RNA probe (Figure 4B). Quantification of the m^6^A RNA-binding affinities shows that they are similar for probes I and IV, and slightly higher for probe III (Figure 4C, Suppl. Figures 4A and 4B). Hence, the content in ribonucleotide U, identified as potentially promoting ECT2 m^6^A-dependent binding to RNA [19], only modestly influence *in vitro* the binding affinity of ECT2 for m^6^A-methylated RNA, and not the ability to form m^6^A-methylated RNA-protein complexes that resist to gel migration. Quite unexpectedly, the EMSAs performed with the non-methylated probes repeatedly showed that the ECT2-Fl protein and its truncated versions 108 and 204 now form stable complexes, albeit with low affinity, when the non-methylated probe lacks U residues (Figure 4A and B, compare the three non-methylated probes). The formation of m^6^A-independent complexes is lost when domain “204 to 302” is deleted (EMSAs with ECT2-303). Finally, as already shown for the smaller constructs ECT2-376 and ECT2-424, no stable complex was observed irrespective of the probe sequence or m^6^A-methylation status (Suppl. Figure 4C). These results show that the ribonucleotide content around the DRACH motif mostly influences the m^6^A-independent binding of ECT2 to RNA, with the presence of uridines that seemingly reduces this m^6^A-independent binding and, as a consequence, might disfavor ECT2 binding to its *in vivo* mRNA targets when they are not m^6^A-methylated.

**Figure 4.**
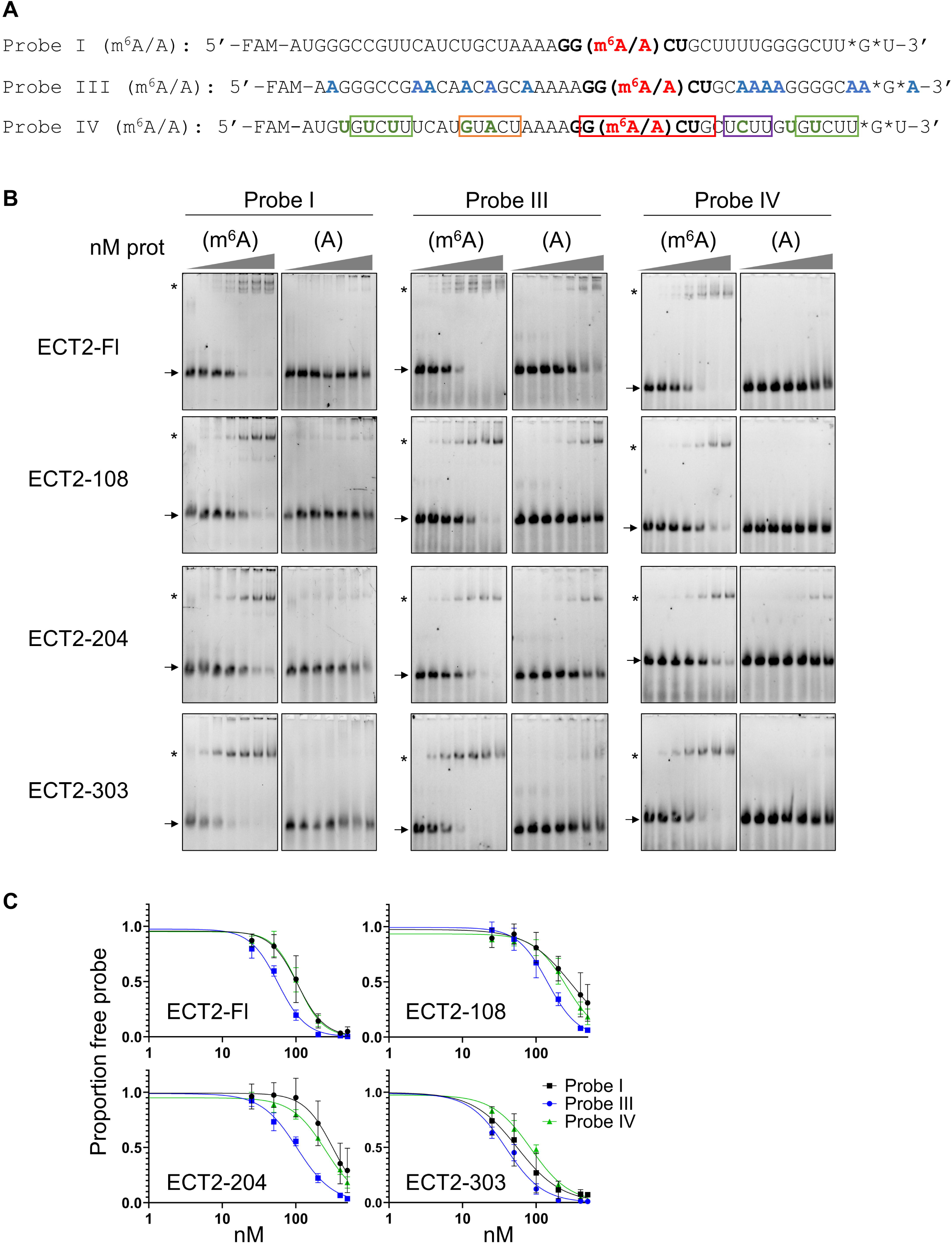
A U-rich ribonucleotide content in the sequence around the DRACH motif reduces m^6^A-independent ECT2 binding. **A.** Sequence of the RNA probes I, III and IV. Each probe is labelled in 5’ with the FAM6 fluorophore. The asterisks indicate thiol-protected bases to avoid degradation of the RNA. The DRACH motif is shown in bold with the methylated or non-methylated adenosine. Probe III was designed from probe I by replacing uridines with adenosines (except in the DRACH motif). Probe IV was also designed from probe I by inserting the motifs described in the work by Arribas-Hernandez and colleagues [19]. This probe contains two UNUNU motifs (in green), a URURU motif (in orange), a YYYY motif (in violet) and the DRACH motif (in red and bold). **B.** EMSA gels obtained with His-SUMO tagged full-length and truncated ECT2 proteins. In each assay, 10 nM of methylated (m^6^A) or non-methylated (A) probes were incubated with increasing concentrations of purified proteins (from 0 to 500 nM). Different probes were used as indicated. Note that EMSA gels shown for probe I are the same in Figure 3B and 4B. The arrows indicate the free probe, and the stars the probe/protein complexes. **C.** Signals of the m^6^A free probes were quantified with the ImageLab software. Graphs show the mean and standard deviation of at least three independent experiments. The curve fittings were obtained using the PRISM software. Each graph allows to compare the curve fittings obtained for one protein and the three different 42 nt methylated RNA probes.

### Structural elements next to ECT2 YTH domain impact on its binding to m^6^A

The YTH domain was first defined more than twenty years ago based on peptide sequence alignment homology search [6]. The subsequent 3D structures of YTH domains demonstrated that evolutionarily distant YTH domain sequences share the same 3D structure that consists in a β-barrel formed by three α helices and six β strands (β1α1β2α2β3β4β5β6α3), defined as the core YTH domain, and revealed the presence of upstream and downstream additional α helices, named α0 and α4, that are present in some but not all of the studied YTH domains. Our previous work based on secondary structure prediction suggested the presence of α0 and α4 helices next to the YTH domains of the angiosperm YTHDF proteins [5]. Alphafold modelling [23] further supports the putative existence of these additional structural elements in ECT2 (Suppl. Figure 5). It predicts a 3_10_ helix for the α0 element (D419-D427), a second 3_10_ helix (D376-R386) located N-terminally of α0, and a very long α helix at the C-terminus of the ECT2 protein (D591-A627) (Suppl. Figure 5A). The first 3_10_ helix which locates some 70 amino-acids upstream of the YTH domain, is well conserved at the primary sequence level amongst the YTHDFA proteins from angiosperms (Suppl. Figures 2A and 5A, in magenta). We named this 10 amino-acid long region the SH domain (for Small Helix). The second 3_10_ helix, named α0 (Suppl. Figure 5A, in blue) and the C*-*terminal helix (in beige) are reminiscent of the α0 and α4 helices found in the extended YTH domain of the human YTHDF1 [8]. Furthermore, Alphafold predicts the existence of several hydrogen bonds between the α4/C-terminal helix and the α0 or SH domain, and between the core YTH and the SH domains or α4/C-terminal helix (Suppl. Figures 5B, 5C), further suggesting that these structural elements spatially locate in close proximity to each other and to the ECT2 YTH domain.

**Figure 5.**
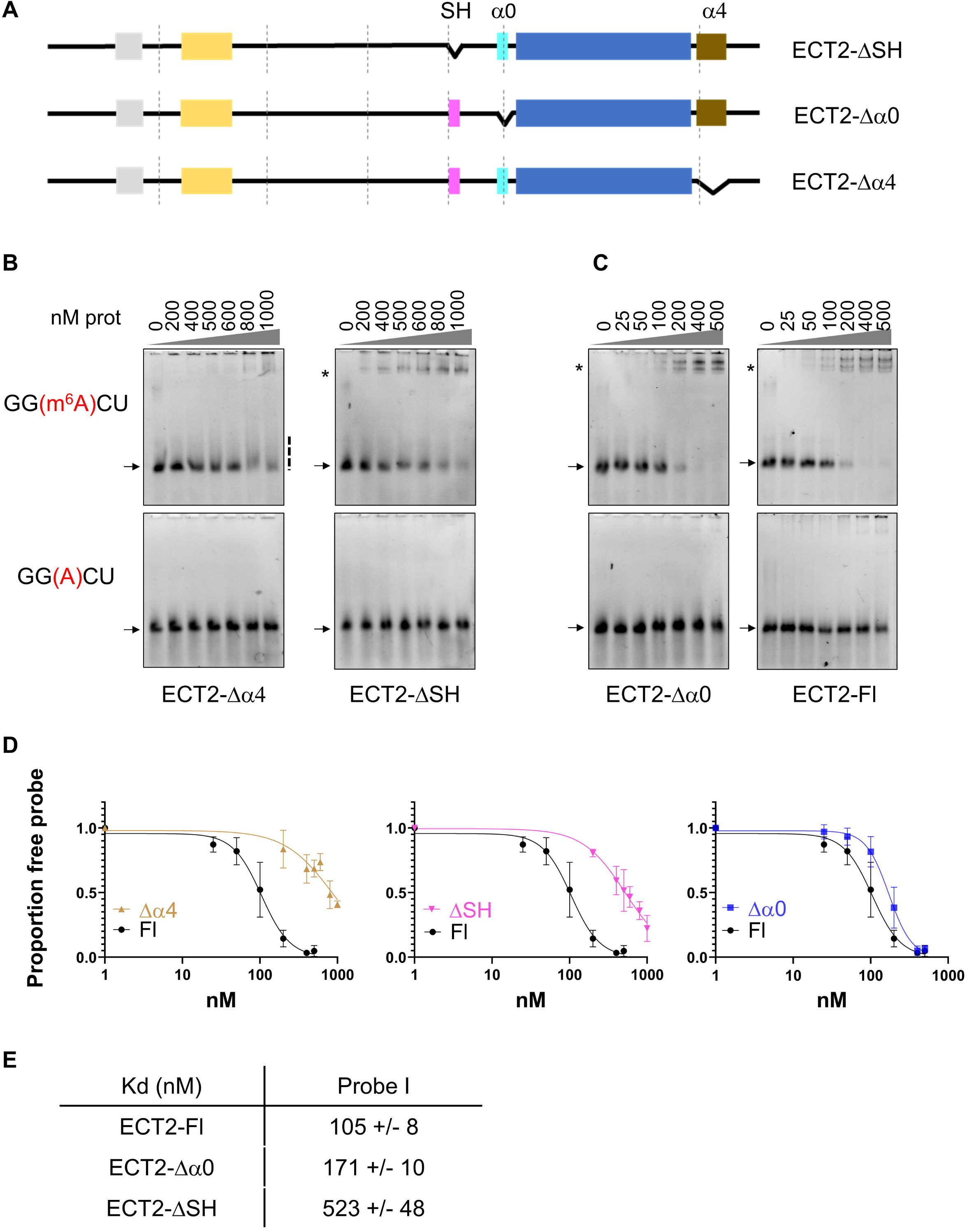
Structural elements next to ECT2 YTH domain impact on its binding to m^6^A. **A.** Graphical representation of the ECT2 deletion mutants. The three structural elements detected through Alphafold are represented in addition to the YTH domain: SH (3_10_ helix, aa 376-386) in magenta, α0 helix (3_10_ helix, aa 419-427) in blue and α4/C-terminal helix (α helix, aa 591-6267) in brown. **B-C.** EMSA gels obtained with the different His-SUMO tagged deletion mutants and the His-SUMO tagged full-length ECT2 (ECT2-Fl). Each purified protein was incubated with 10 nM of methylated RNA probe (probe I, see Figure 3A). The protein concentration ranged from 0 to 1000 nM for the ECT2-Δα4 and ECT2-ΔSH mutants (B), and from 0 to 500 nM for the ECT2-Δα0 mutant and ECT2-Fl (C). The arrows and stars respectively indicate the free probe and the probe/protein complexes, and the dashed line, the smear observed when unstable complex. **D-E.** Quantification of the signals of the m^6^A free probes using the ImageLab software. Graphs show the mean and standard deviation of three independent experiments. The fitting curves (D) and apparent Kds (E) were obtained using the PRISM software.

Based on these observations, we hypothesized that one or more of these elements might contribute to ECT2 YTH domain binding to m^6^A-RNA. We therefore produced recombinant ECT2 proteins lacking either the SH domain, α0 or α4 helices (Figure 5A, Suppl. Figure 5D) and performed EMSAs. Deletion of the α4 helix or the SH domain triggers a clear decrease in binding affinity to m^6^A-RNA (Figure 5B, 5D). For the ΔSH (D376-386) mutant, a complex with m^6^A-methylated RNA is still visible conversely to the Δ α4 mutant (Figure 5B and 5D-E, note that here the range of protein concentration used is higher than in other EMSAs). The Δα0 (Δ419-427) mutant mostly behaves as the full-length ECT2 protein (Figure 5C), with only a slight difference in the affinities (Figures 5D right panel, and 5E). Of note, deletion of α0 does not affect the formation of an RNA-protein complex that migrates as a doublet, while loss of the SH domain does, suggesting that formation of a putative ECT2 dimer not only relies on the “1-108” domain but also on the SH domain. In summary, two domains outside the IDR, the α4 helix and the SH domain, also modulate ECT2 m^6^A-dependent binding to RNA *in vitro*.

## Discussion

Biophysical and biochemical studies conducted with the YTH domains of human YTHDF and YTHDC proteins, rat YTHDC1, yeast Mrb1 and Pho92 and Arabidopsis CPSF30-L proteins indicated that YTH domains can be self-sufficient to recognize the m^6^A residue and bind m^6^A-methylated RNA. Here we show that Arabidopsis ECT2 YTH domain alone poorly binds to m^6^A *in vitro*, in comparison to human YTHDC1 and Arabidopsis CPSF30-L YTH domains and ECT2 full-length protein. Importantly, we identify regions in ECT2 N-terminal IDR as well as structural elements located in close proximity to its YTH domain that enhance its binding to m^6^A (see summary of finding in Suppl. Figure 6). Below, we discuss these findings in the light of the recent *in vivo* studies on ECT2, which proposed that ECT2 N-terminal IDR may regulate its binding to m^6^A-methylated mRNAs. Together these *in vitro* and *in vivo* findings open new perspectives on the regulation of YTH domain binding to m^6^A and its biological relevance.

Every YTHDF proteins from angiosperms carry upstream of their YTH domain a long IDR, which in the case of ECT2 contact the m^6^A-methylated target mRNAs *in vivo*, at U-rich motifs, located next to the methylated DRACH motif. The authors suggested that these contacts between the IDR and the RNA may stabilize the reader on the m^6^A residue [19]. Our results not only provide the direct demonstration that ECT2 IDR is key for the formation of a stable m^6^A complex but also map the region located between amino acids 300 and 375, as the putative mediator of this stabilization. We also find that, conversely to the studied YTH readers from mammals and *Z. rouxii* and *S. cerevisiae* [7–10], ECT2 is not able to form a complex with a short 11 nt-long m^6^A-methylated RNA probe. This observation is again fully consistent with the *in vivo* data supporting that the ECT2 IDR directly binds RNA targets next to the m^6^A site [19]. Our work also suggests that ECT2 IDR may carry additional regulatory regions within the domains “1-107” and “204-302”, that would respectively increase or decrease the binding affinity to the m^6^A RNA probe. Such RNA binding regulatory function is one of the reported properties of IDRs connecting or surrounding RNA Binding Domains [24]. Considering that ECT2 IDR has been consistently found to be phosphorylated at several positions, in proteome- wide studies [26], a tempting hypothesis is that phosphorylation, and possibly other post- translational modifications, may regulate its YTH domain binding to m^6^A.

Our data also show that the uridine content around the DRACH motif plays a significant role in modulating the affinity of ECT2 to the non-methylated form of the RNA. Based on this finding, together with the *in vivo* iCLIP results showing that ECT2 IDR also contacts the m^6^A- methylated mRNA targets in close proximity to the DRACH motif [19], we propose a model where the combination of ECT2 YTH domain and ECT2 IDR bindings to RNA further increases the selectivity of ECT2 towards the m^6^A-methylated version of its mRNA targets. *In planta*, ECT2 target RNAs are enriched in uridines in the vicinity of the DRACH motif, while *in vitro* ECT2 binds preferentially RNAs with a poor uridine content, but only when they are non-methylated. Taken together these results indicate that ECT2 *in vivo* has a proprension for not interacting with RNA sequences enriched in uridines, unless they possess a m^6^A-methylated DRACH motif.

From *in vivo* studies, two domains of the ECT2 N-terminus, tagged as N3.2/SLIM (aa 48-86) and N8 (aa 355-394), are required for its function in leaf emergence [20]. Interestingly, our study indicates that the N8 domain is potentially important for ECT2’s m^6^A RNA binding activity, since N8 covers the 20 last residues of region 303-376 plus the SH domain, which are both required for higher affinity binding *in vitro*. Our data also explain why N8 and N3.2 have additive physiological roles, with N8 that would mediate m^6^A-methylated RNA interaction while N3.2/SLIM mediates ECT2 interaction with a poly(A) binding protein [20].

In addition to ECT2 IDR, we report on the importance of structural elements close to its YTH domain, some of which, like the α0 and α4 helices, have already been identified in other YTH proteins. Alphafold predictions from ECT2 and sequence comparisons, strongly suggest the presence of the α0 and α4 helices, that are located, respectively on the N- and C-terminal sides of the YTH sequence, as well as a previously unrecognized 3_10_ helix (aa 376-386) that we named SH for Small Helix, located on the N-terminal part of ECT2’s YTH sequence upstream of α0. These structural elements are not sufficient for ECT2 YTH domain to form a stable complex with m^6^A-RNA in our EMSAs, as shown through the use of the ECT2-376 protein, yet both of them are necessary. The α0 helix seems to be dispensable and only has a moderate effect on the affinity regulation, whereas the SH strongly impacts on the affinity and the α4 helix is absolutely necessary for the formation of a m^6^A-RNA protein complex that resists to gel migration. Since the two upstream helices (SH and α0) and the downstream α4 are predicted to share hydrogen bonds and to fold back over ECT2 core YTH domain (Suppl. Figure 5), this suggests that together they may form a unique structural element which contributes to ECT2 binding to m^6^A. Noticeably, *in vivo*, deletion of ECT2 last 56 residues (aa 612-667) does not affect its physiological role in leaf emergence even though part of α4 appears to be deleted [20]. Yet, it cannot be concluded that the partially-truncated α4 helix has no functional importance, since a large portion of the α4 helix (20 residues) remains expressed and this may be enough for its function in ECT2 RNA binding activity. Whatever is the exact *in vivo* relevance of ECT2 α4 helix, our study shows the importance of structural elements that are in close proximity to its YTH domain and that may form together a unique and functionally important structural element.

Amongst flowering plants, the joint presence of SH, α0 and α4 domains appears conserved only in the DFA and DFB clades. While an α0 is predicted to exist in all YTHDF angiosperm proteins [5], only members of the DFA and DFB share a well-conserved SH domain, indicating a possible difference in the mode of binding to m^6^A-methylated RNAs for members of the DFC subclade (Suppl. Figures 2, and 7-8).

On a broader evolutionary perspective, the YTH domain of the fly YTHDC1 protein, seems to behave like ECT2 YTH domain. Indeed, an α0-α4 extended YTH domain of YTHDC1 fails to form a stable complex with an m^6^A-RNA probe as tested by EMSA [12]. In contrast, YTH domains from the yeasts *Z. rouxii* Mrb1 protein and *S. cerevisiae* Pho92 protein, as well as human YTHDC2, lack the α0 and α4 helices (Suppl. Figure 8) yet they are self-sufficient to interact with a short (5-11nt) m^6^A RNA *in vitro*, indicating that these YTH domains must present slight structural differences responsible for their differential capacity to bind, or not bind, m^6^A-methylated RNA in a self-sufficient manner. The human YTHDF1-2-3 and YTHDC1, as well as rat YTHDC1 proteins possess α0 and α4 helices, according to structural studies [8–10] or Alphafold prediction (Suppl. Figure 8). Self-sufficient binding to m^6^A- methylated RNA has been demonstrated but only in the case of their extended YTH domain (which includes the peripheral α0 and α4 helices), and not yet for their core YTH domain only. This is also the case for the Arabidopsis CPSF30-L YTH reader. Whatever the exact contribution of α0 and α4 helices are in these YTH proteins, they illustrate that the existence of regulatory regions in proximity to the YTH domain is not limited to the readers of the flowering plants but might be widespread among eukaryotes. Furthermore, our results also show that regulatory regions might not be limited to the structural elements present in the close proximity of the YTH domain but also present in surrounding IDRs. Knowing that the vast majority of YTH proteins possess at least one large IDR, in addition to the α0 and α4 helices, a sizeable proportion of YTH proteins may possess peptide sequences that regulate the binding of their YTH domains to m^6^A-methylated RNA with *in vivo* relevance that remain to be explored.

## Supplemental Tables

Table S1. List of primers and RNA probes

Table S2. Amino acid sequences of extended YTH domains used for biochemical characterization

Table S3. Amino acid sequences of YTH proteins from Viridiplantae

## Acknowledgements

We thank the EMBL (Heidelberg, Germany) for providing the expression vector pETM11- SUMO3-eGFP, and Ramesh Pillai for providing the expression vector pETM11SS. We thank Jean-Marc Deragon for providing the sequences of the angiosperm YTHDF proteins. We thank Andrès Palencia and Ramesh Pillai for their helpful comments. CBA thanks the “Laboratoires d’Excellences (LABEX)” TULIP (ANR-10-LABX-41) and of the “École Universitaire de Recherche (EUR)” TULIP-GS (ANR-18-EURE-0019), in the frame of which the work conducted at LGDP in Perpignan was set. This work used the platforms of the Grenoble Instruct-ERIC center (ISBG; UAR 3518 CNRS-CEA-UGA-EMBL) within the Grenoble Partnership for Structural Biology (PSB), supported by FRISBI (ANR-10-INSB-0005-02) and GRAL, financed within the University Grenoble Alpes graduate school (Ecoles Universitaires de Recherche) CBH-EUR-GS (ANR-17-EURE-0003). The Institut de Biologie Structurale acknowledges integration into the Interdisciplinary Research Institute of Grenoble. MRJ is laureate of the Impulscience® program of the Fondation Bettencourt Schueller.

## Funding

The study was supported by grant from the French Agence Nationale pour la Recherche (ANR) “Heat-EpiRNA” (ANR-17-CE20-007-01) to CBA and AV.

## Author contributions

DSB, CBA and AV conceived and designed the experiments. DSB, CK, KY, CGL, JJF did the cloning. DSB, CK, KY did the production of proteins in *E. coli*. DSB did the protein purifications and the EMSAs. ED, MRJ acquired the ^1^H NMR spectrum.

DSB, CBA and AV analyzed the data, and wrote the paper. All authors reviewed the results and approved the final version of the manuscript.

## Data availability statement

Data sharing is not applicable to this article as no datasets were generated or analyzed during the current study

## Notes

### Competing Interest Statement

The authors have declared no competing interest.

