## Supplemental Figures for "ECT2 peptide sequences outside the YTH domain regulate its m^6^A-RNA binding"

#### Supplemental Figure 1

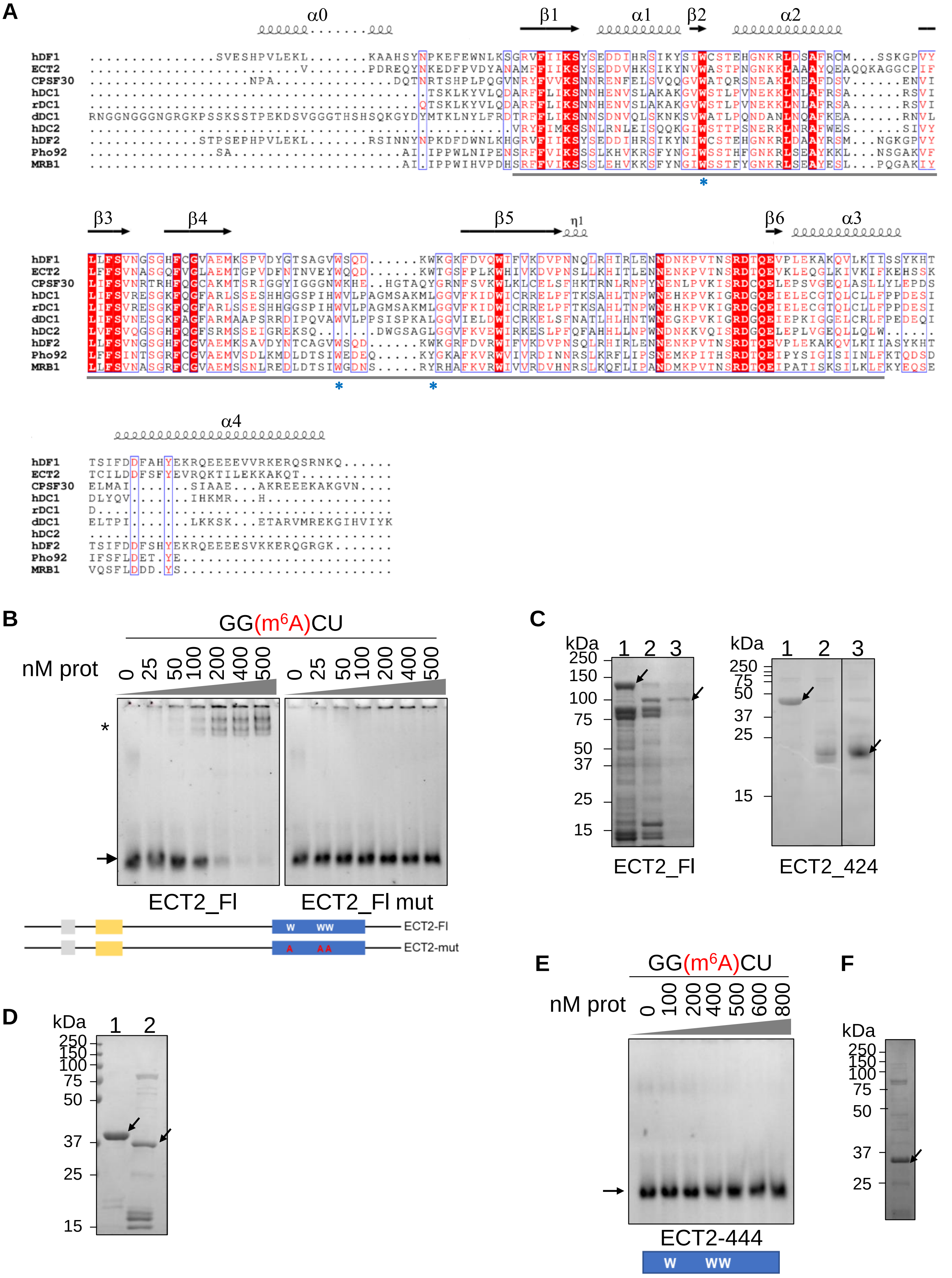

#### Supplemental Figure 2

**A**

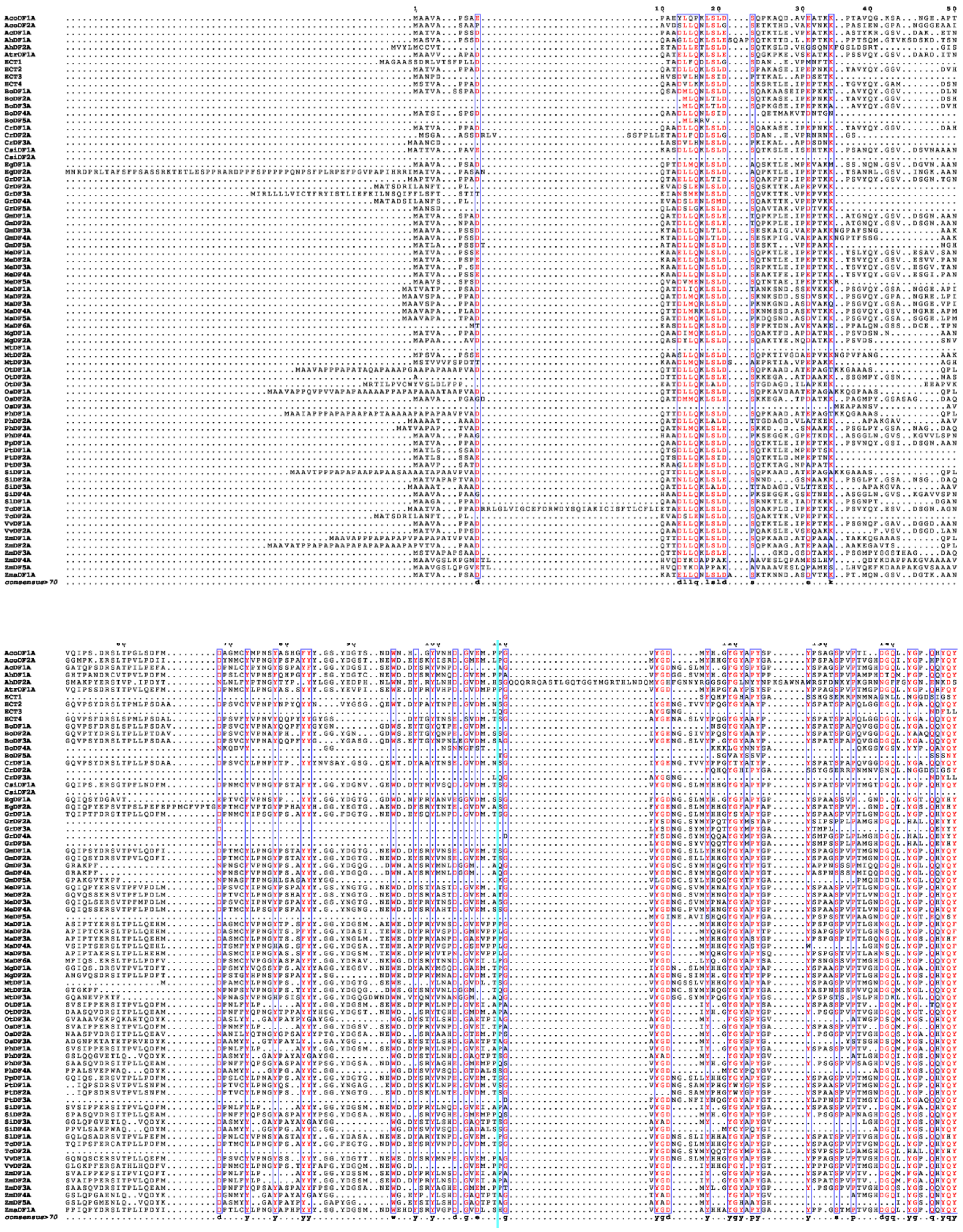

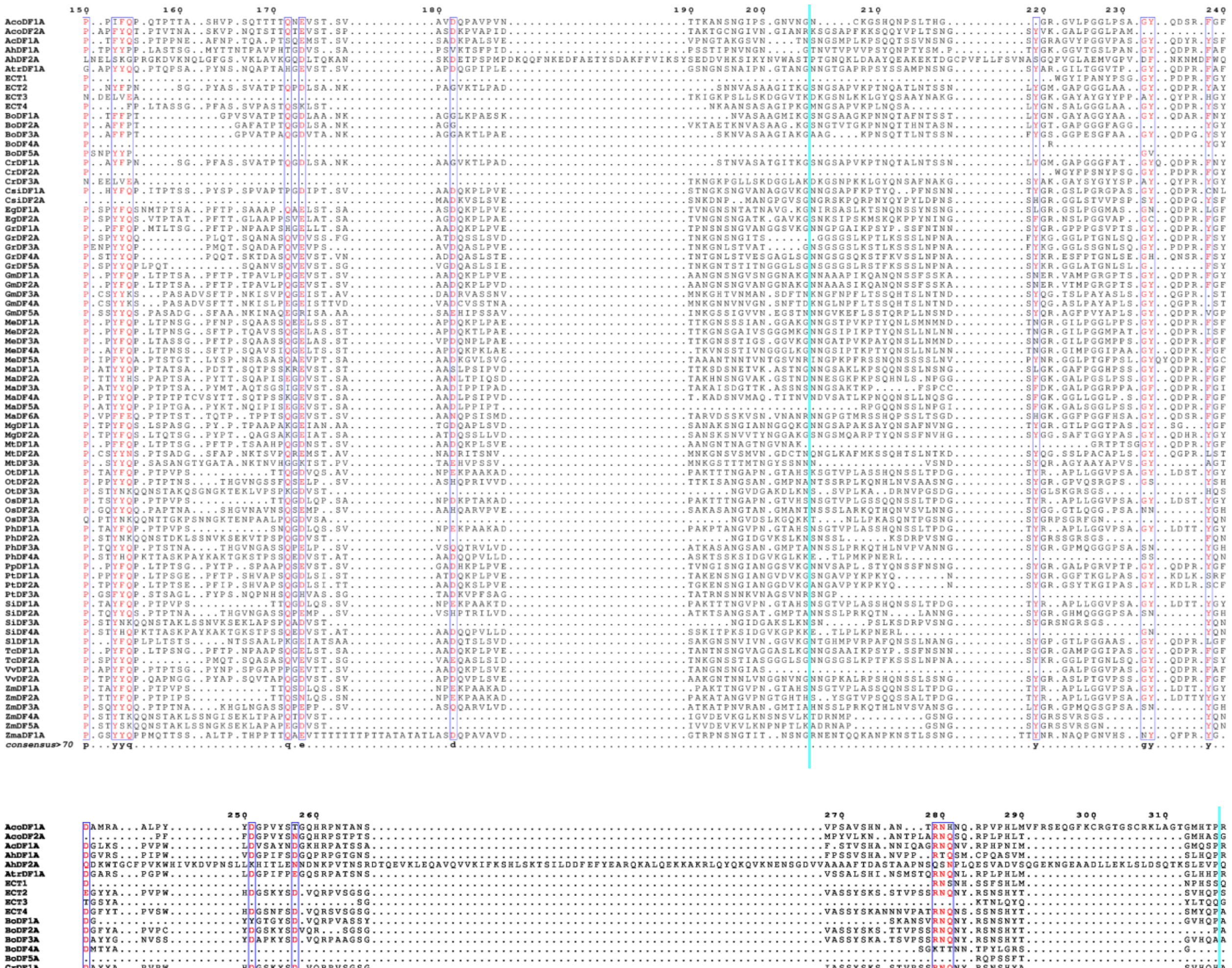

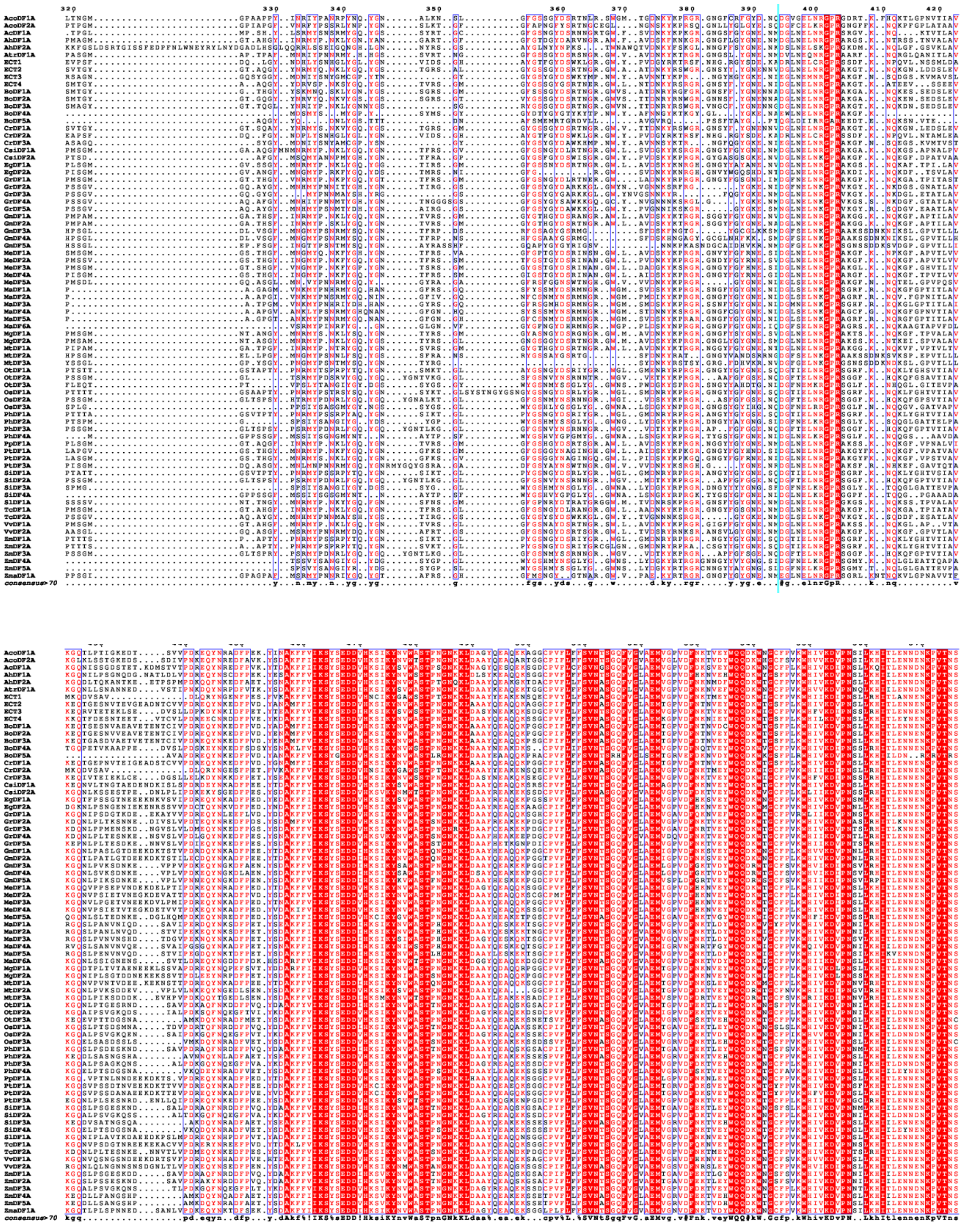

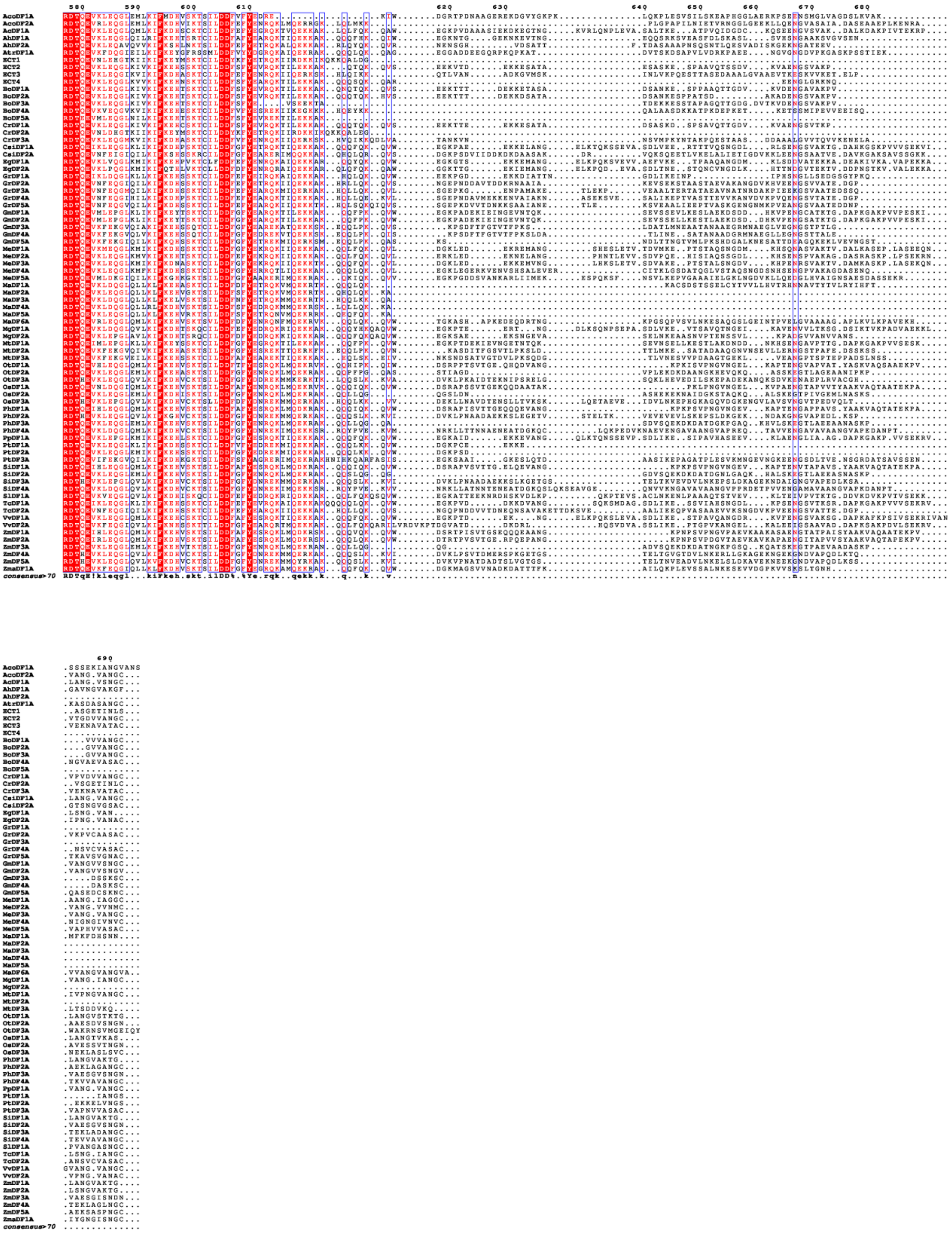

#### Supplemental Figure 2

**B**

```
MATVAPPADQATDLLQKLSLDSPAKASEIPEPNKKTAVYQYGGVDVHGQVPSYDRSLTPMLPSDAADPSVC
YVPNPYNPYQYYNVYGSGQEWTDYPAYTNPEGVDMNSGIYGENGTVVYPQGYGYAAYPYSPATSPAPQLGG
EGQLYGAQQYQYPNYFPNSGPYASSVATPTQPDLSANKPAGVKTLPADSNNVASAAGITKGSNGSAPVKPT
NQATLNTSSNLYGMGAPGGGLAAGYQDPRYAYEGYYAPVPWHDGSKYSDVQRPVSGSGVASSYSKSSTVPS
SRNQNYRSNSHYTSVHQPSSVTGYGTAQGYYNRMYQNKLYGQYGSTGRSALGYGSSGYDSRTNGRGWAATD
NKYRSWGRGNSYYYGNENNVDGLNELNRGPRAKGTKNQKGNLDDSLEVKEQTGESNVTEVGEADNTCVVPD
REQYNKEDFPVDYANAMFFIIKSYSEDDVHKSIKYNVWASTPNGNKKLAAAYQEAQQKAGGCPIFLFFSVN
ASGQFVGLAEMTGPVDFNTNVEYWQQDKWTGSFPLKWHIVKDVPNSLLKHITLENNENKPVTNSRDTQEVK
LEQGLKIVKIFKEHSSKTCILDDFSFYEVRQKTILEKKAKQTQKQVSEEKVTDEKKESATAESASKESPAA
VQTSSDVKVAENGSVAKPVTGDVVANGC-
```

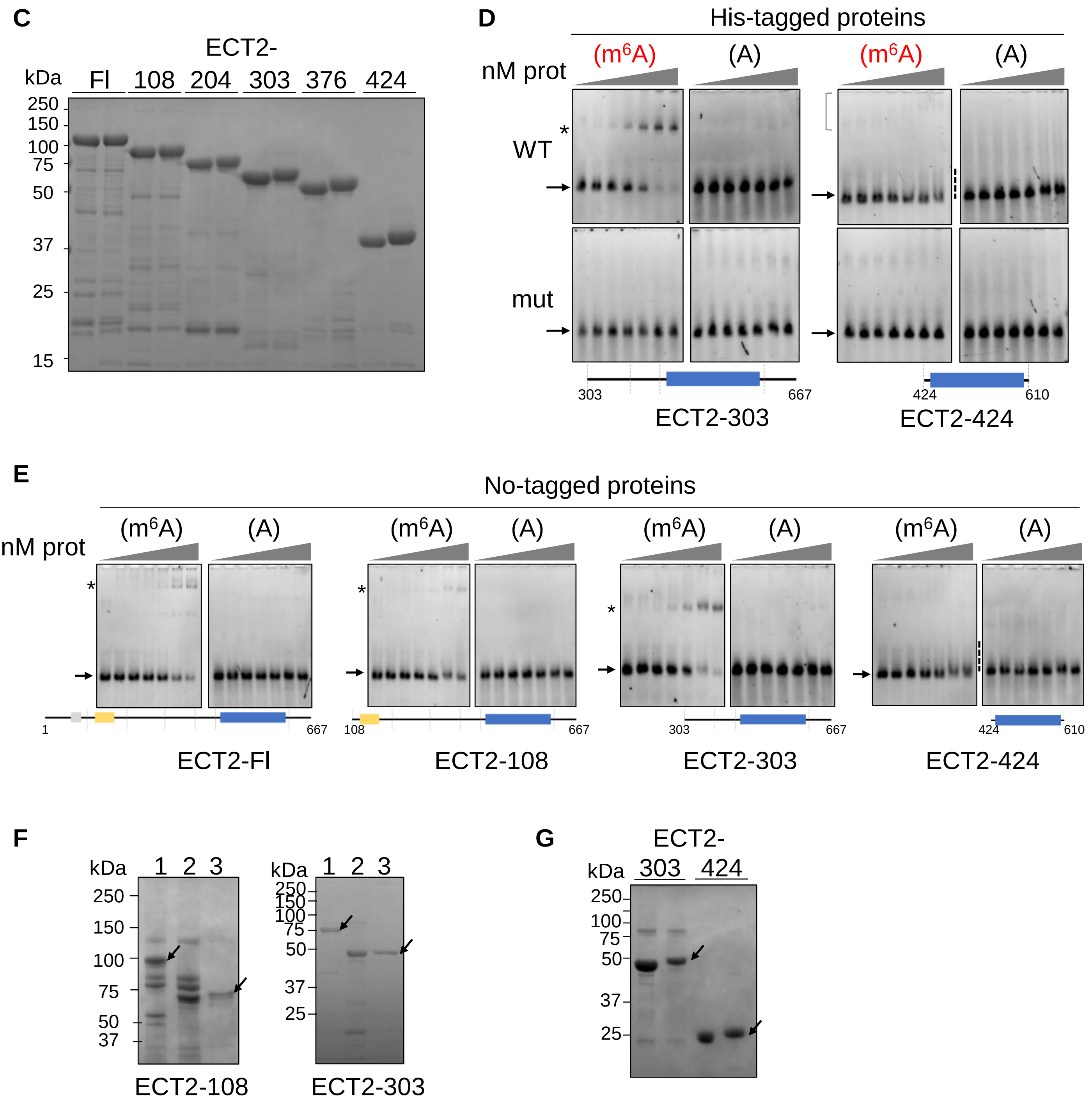

**Supplemental Figure 3**

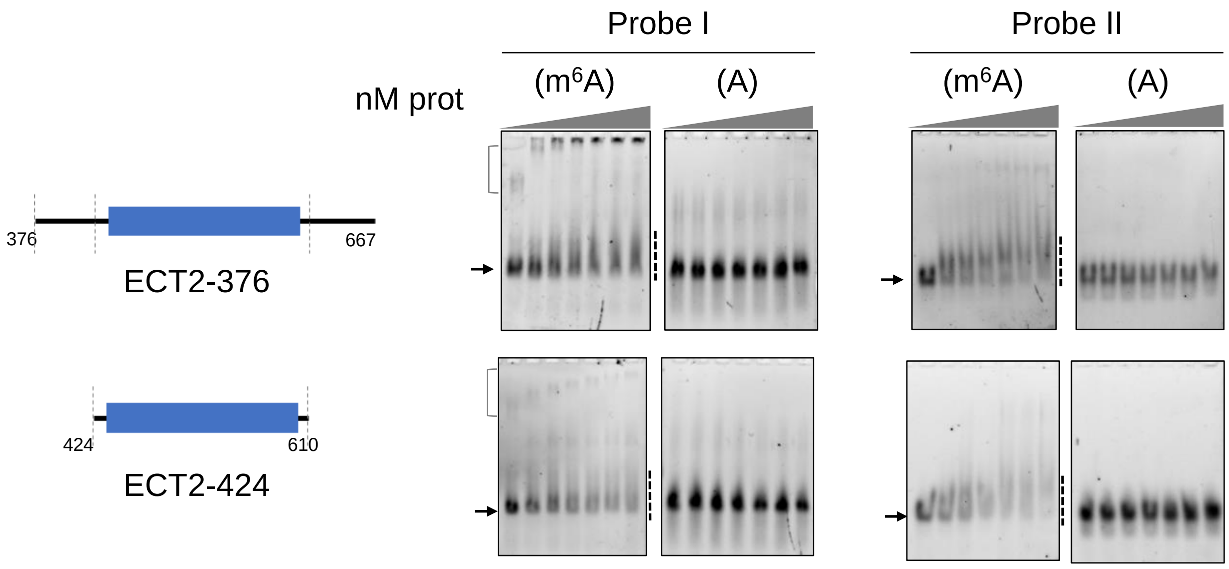

#### Supplemental Figure 4

**A**

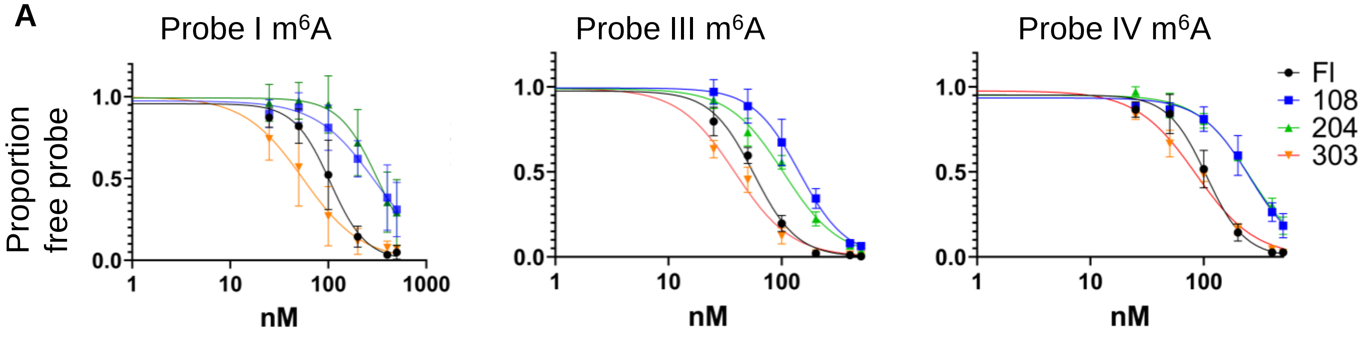


```
Probe I m6A:   5'-FAM-AUGGGCCGUUCAUCUGCUAAAAGG(m6A)CUGCUUUUGGGGCUU*G*U-3'
Probe III m6A: 5'-FAM-AAGGGCCGAACAACAGCAAAAAGG(m6A)CUGCAAAAGGGGCAA*G*A-3'
Probe IV m6A:  5'-FAM-AUGUGUCUUUCAUGUACUAAAAGG(m6A)CUGCUCUUGUGUCUU*G*U-3'
```

**B**

| Kd (nM) | Probe I m$^6$A | Probe III m$^6$A | Probe IV m$^6$A |
|---|---|---|---|
| ECT2 FI | 105 +/- 8 | 57 +/- 3 | 106 +/- 7 |
| ECT2-108 | 296 +/- 43 | 143 +/- 18 | 255 +/- 18 |
| ECT2-204 | 318 +/- 35 | 104 +/- 6 | 253 +/- 15 |
| ECT2-303 | 55 +/- 7 | 39 +/- 2 | 85 +/- 5 |

**C**

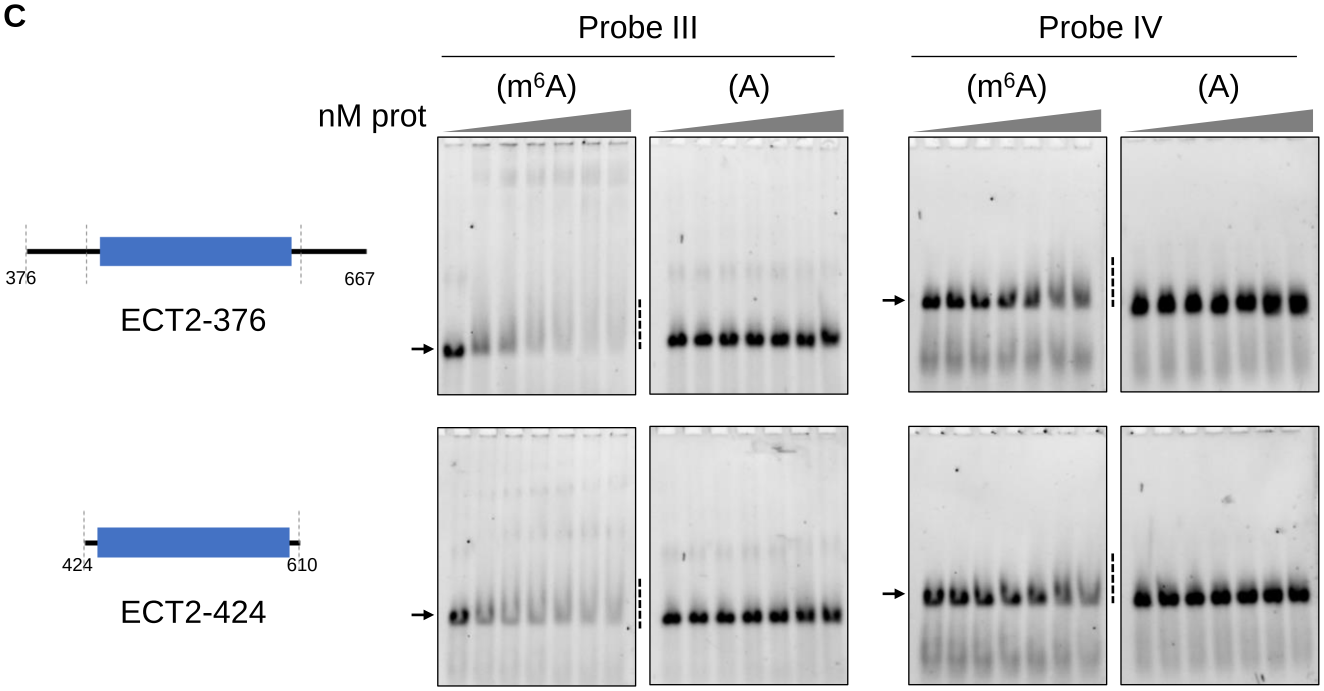

**Supplemental Figure 5**

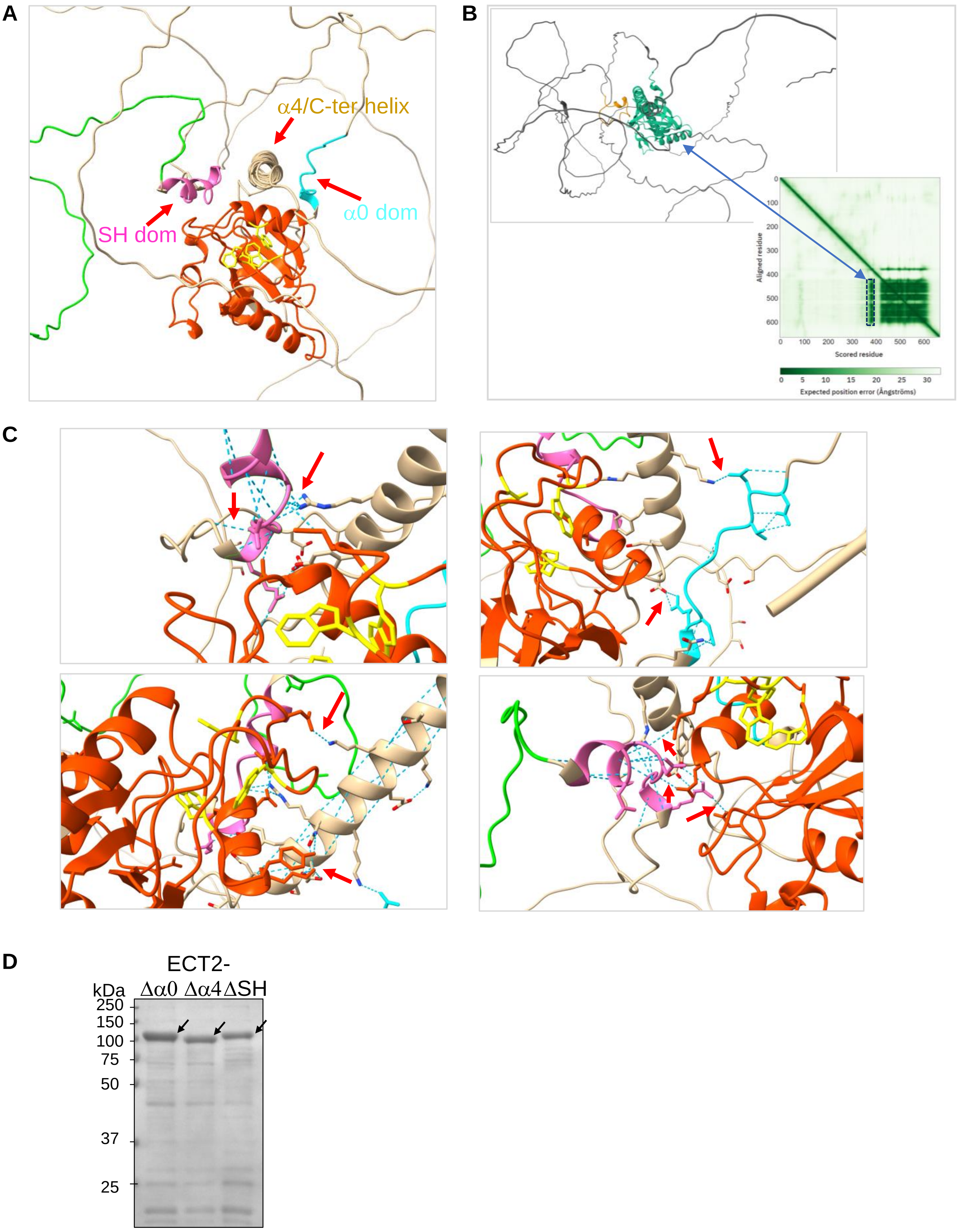

### Supplemental Figure 6

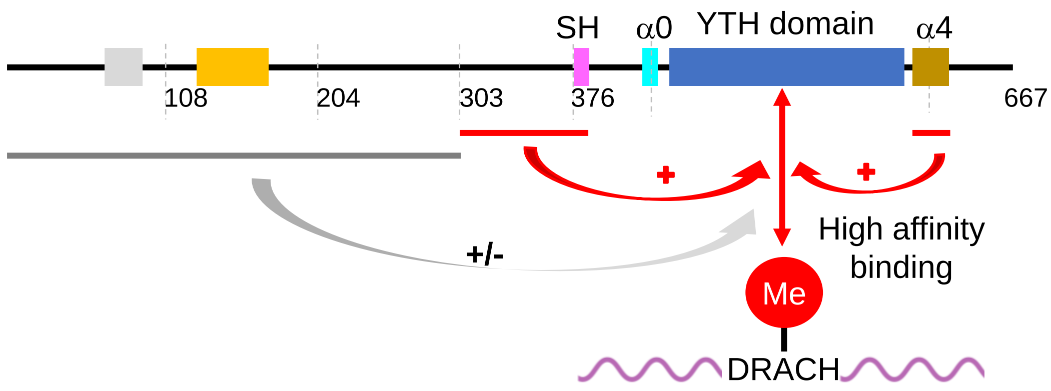

### Supplemental Figure 7

DFB

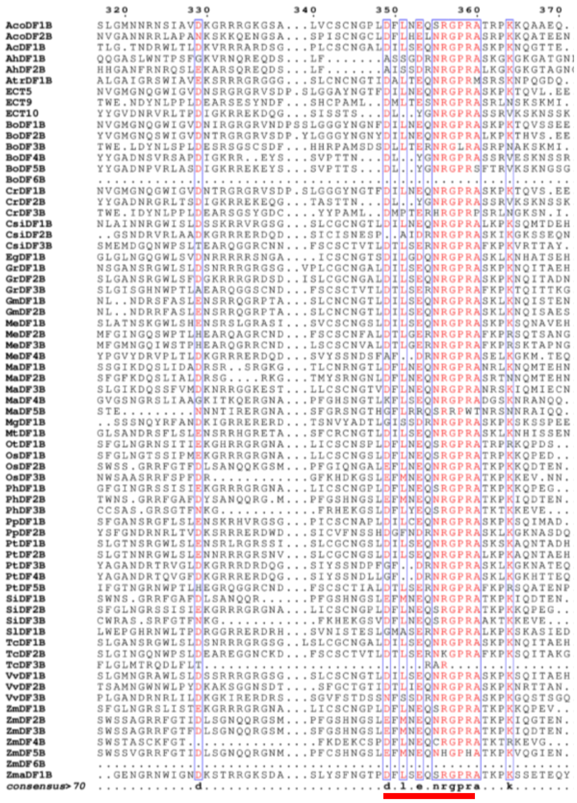

DFC

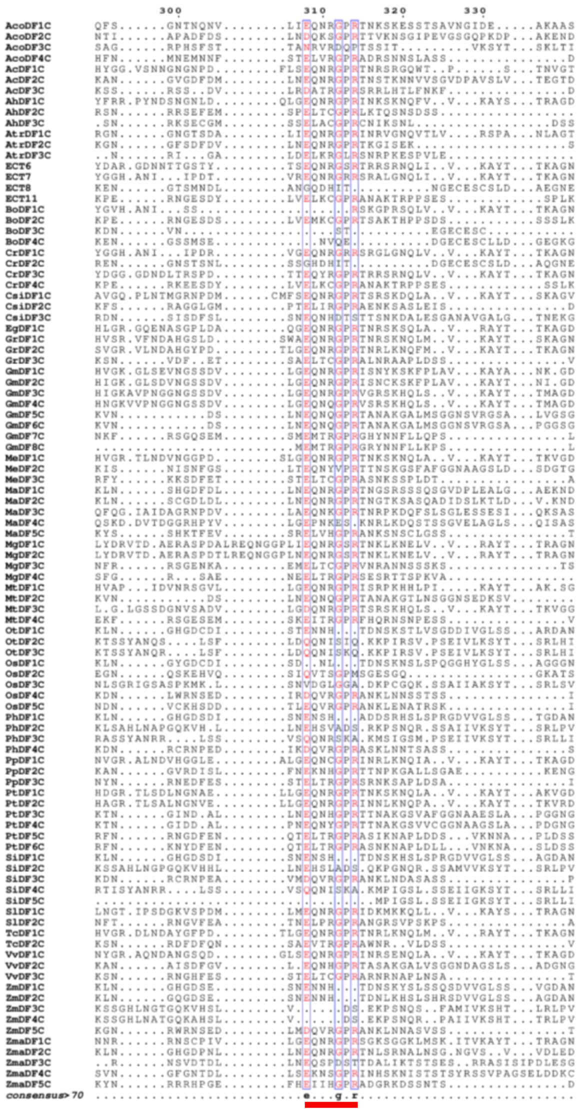

#### Supplemental Figure 8

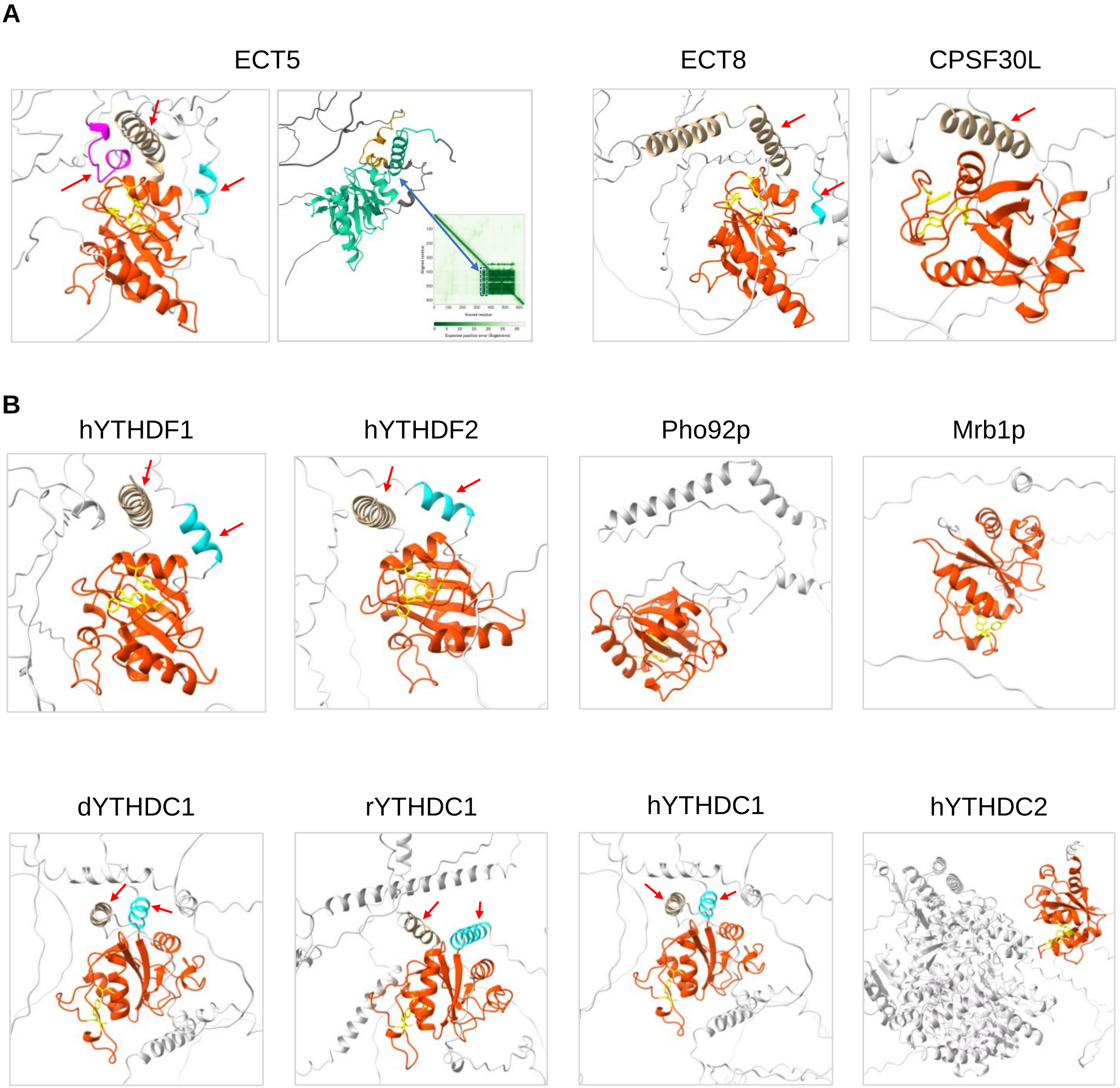
